# High-affinity, neutralizing antibodies to SARS-CoV-2 can be made in the absence of T follicular helper cells

**DOI:** 10.1101/2021.06.10.447982

**Authors:** Jennifer S. Chen, Ryan D. Chow, Eric Song, Tianyang Mao, Benjamin Israelow, Kathy Kamath, Joel Bozekowski, Winston A. Haynes, Renata B. Filler, Bridget L. Menasche, Jin Wei, Mia Madel Alfajaro, Wenzhi Song, Lei Peng, Lauren Carter, Jason S. Weinstein, Uthaman Gowthaman, Sidi Chen, Joe Craft, John C. Shon, Akiko Iwasaki, Craig B. Wilen, Stephanie C. Eisenbarth

## Abstract

T follicular helper (Tfh) cells are the conventional drivers of protective, germinal center (GC)-based antiviral antibody responses. However, loss of Tfh cells and GCs has been observed in patients with severe COVID-19. As T cell-B cell interactions and immunoglobulin class switching still occur in these patients, non-canonical pathways of antibody production may be operative during SARS-CoV-2 infection. We found that both Tfh-dependent and -independent antibodies were induced against SARS-CoV-2 as well as influenza A virus. Tfh-independent responses were mediated by a population we call lymph node (LN)-Th1 cells, which remain in the LN and interact with B cells outside of GCs to promote high-affinity but broad-spectrum antibodies. Strikingly, antibodies generated in the presence and absence of Tfh cells displayed similar neutralization potency against homologous SARS-CoV-2 as well as the B.1.351 variant of concern. These data support a new paradigm for the induction of B cell responses during viral infection that enables effective, neutralizing antibody production to complement traditional GCs and even compensate for GCs damaged by viral inflammation.

**One-Sentence Summary:** Complementary pathways of antibody production mediate neutralizing responses to SARS-CoV-2.

## Main Text

Antibodies are critical for protection against severe acute respiratory syndrome coronavirus 2 (SARS-CoV-2), the causative agent of the coronavirus disease 2019 (COVID-19) pandemic (*1*). T follicular helper (Tfh) cells are the conventional drivers of protective antibody responses, as they support immunoglobulin class switching, germinal center (GC)-based affinity maturation, and long-lived humoral immunity (*2*). Indeed, multiple studies have reported a correlation between circulating Tfh (cTfh) cells – in particular, type 1 CXCR3^+^CCR6^−^ cTfh cells – and neutralizing antibody titers in COVID-19 patients (*3–9*). While there have been mixed findings on the association of Tfh cells with disease severity, several groups have observed an absence of Tfh cells and GCs in severely ill patients (*4, 5, 7, 9–15*). Despite a loss of Tfh cells and GC structures, T cell-B cell interactions and antibody class switching still occur in the secondary lymphoid organs of these patients (*14*). Findings of enhanced extrafollicular B cell responses associated with severe disease further suggest that non-canonical pathways of antibody production may be operative in these individuals (*14, 16*). However, a causal relationship between antibody provenance and disease severity cannot be established by existing human studies. Thus, it remains unclear whether antibodies produced through non-canonical pathways without Tfh cell help are also protective against SARS-CoV-2.

No study has mechanistically evaluated the role of Tfh cells in antibody production to SARS-CoV-2. Previous work with SARS-CoV has shown that CD4^+^ T cells are required for neutralizing antibody titers (*17*), but the nature of the T cell population has not been identified. For other viruses, as well as bacteria, Tfh cells are thought to be required for class-switched, pathogen-specific antibody production, especially at later timepoints (*18–26*). In contrast, different studies have found conflicting results on the requirement of Tfh cells during vaccination in mice; certain vaccine strategies induce robust Tfh-independent antibody responses, though of lower affinity (*27, 28*), while other strategies fail to induce durable, class-switched, or high-affinity antibodies in mice with Tfh cell deficiency or dysfunction (*19, 29–31*). In humans though, cTfh cell populations in the blood correlate with the response to vaccination (*32–34*). Taken together, while non-Tfh CD4^+^ T cells can promote effective antibodies in certain contexts, protective anti-pathogen humoral immunity is largely thought to be Tfh cell-dependent.

We therefore directly tested whether non-Tfh CD4^+^ T cells could compensate for Tfh cell loss in severe COVID-19, and perhaps during acute viral infection in general, by promoting class-switched antibodies. Based on prior work we hypothesized that antibodies induced through such non-canonical mechanisms would be lower in quantity and quality. To test this, we characterized the titer, isotype, longevity, affinity, and function of antibodies to SARS-CoV-2 as well as another clinically relevant respiratory virus, influenza A virus, from Tfh-deficient mice. We found that both infections induced substantial levels of Tfh-independent class-switched antibodies, likely driven by a population we call lymph node (LN)-Th1 cells. LN-Th1-driven antibodies to SARS-CoV-2 were durable and remarkably high-affinity. These antibodies were also broadly reactive against diverse SARS-CoV-2 epitopes and effective at neutralizing both homologous and heterologous viruses. In addition, LN-Th1 cells were present in the setting of intact GCs, suggesting that LN-Th1-driven responses are not just compensatory but also complementary with canonical Tfh-dependent antibody production to multiple respiratory viruses. Thus, our study suggests a new paradigm for T cell-driven antibody responses to viruses in which a subset of Th1 cells complement Tfh cells in secondary lymphoid organs by interacting with and guiding B cell responses without emigrating to tissues to direct local cellular responses. Therefore, multiple types of CD4^+^ T cells in lymphoid organs coordinate the humoral response to viruses. This new understanding of T cell-B cell interaction could aid in approaches to develop effective protection against SARS-CoV-2 as well as other viruses.

### Respiratory viral infections induce Tfh-dependent and -independent antibodies

To study the cellular pathways that promote antibody production to SARS-CoV-2, we used mice that lack different CD4^+^ T cell subsets. *Bcl6*^fl/fl^*Cd4*^Cre^ mice lack Tfh cells due to deletion of the Tfh lineage-defining transcription factor BCL6, and *Ciita*^−/−^ mice lack all CD4^+^ T cells due to loss of MHC class II expression (*30, 35*). As SARS-CoV-2 is unable to efficiently interact with mouse ACE2, we overexpressed human ACE2 (hACE2) in the respiratory tract of these mice via intratracheal administration of AAV-hACE2 (*36, 37*). Two weeks after AAV-hACE2 transduction, we infected mice intranasally with SARS-CoV-2 (isolate USA-WA1/2020) and then assessed cellular and humoral responses at 14 days post infection (dpi) (Fig. 1A). PD-1^hi^CXCR5^hi^ Tfh cells were efficiently deleted in the mediastinal LN (medLN) of *Bcl6*^fl/fl^*Cd4*^Cre^ mice (Fig. S1, A to C). Consistent with the loss of Tfh cells, GC B cells were severely impaired, and plasmablast formation was reduced (Fig. S1, D to H). Yet viral burden in the lungs of *Bcl6*^fl/fl^ and *Bcl6*^fl/fl^*Cd4*^Cre^ mice was similar at 7 dpi (Fig. S1I).

**Figure 1:**
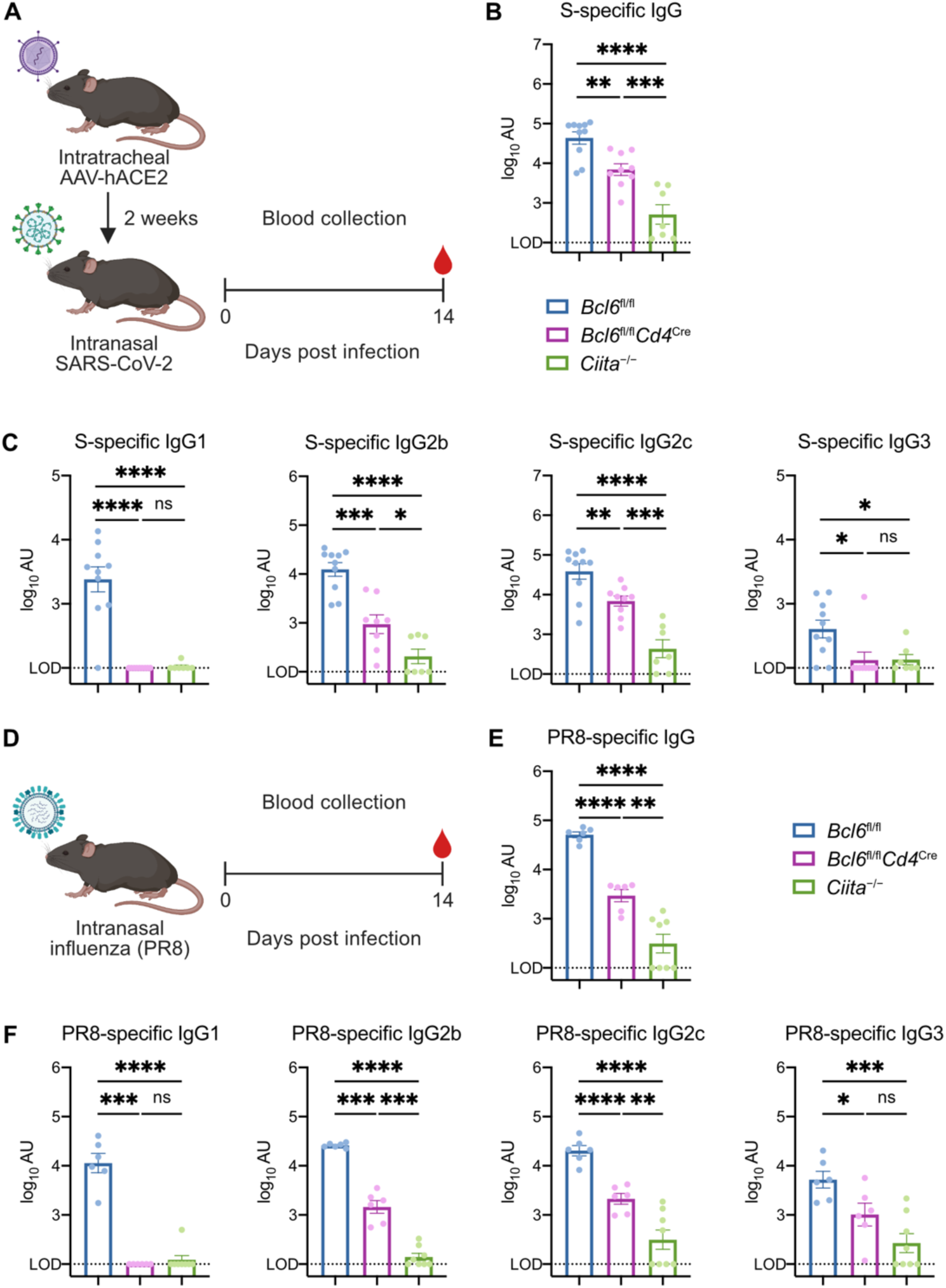
Respiratory viral infections induce Tfh-dependent and -independent antibodies. (**A**) Schematic of experimental design for assessing serum antibody responses against SARS-CoV-2. Mice were infected intranasally with SARS-CoV-2 (isolate USA-WA1/2020) two weeks following intratracheal administration of AAV-hACE2. Sera were collected 14 days post infection (dpi) for quantification of antibody titers by ELISA. (**B**) Spike (S)-specific total IgG antibody titers in sera from control (*Bcl6*^fl/fl^; blue symbol), Tfh-deficient (*Bcl6*^fl/fl^*Cd4*^Cre^; magenta symbol), or CD4^+^ T cell-deficient (*Ciita*^−/−^; green symbol) mice at 14 dpi with SARS-CoV-2. (**C**) S-specific IgG1, IgG2b, IgG2c, and IgG3 antibody titers at 14 dpi with SARS-CoV-2. (**D**) Schematic of experimental design for assessing serum antibody responses against influenza. Mice were infected with mouse-adapted influenza virus (PR8) by intranasal administration. Sera were collected at 14 dpi for quantification of antibody titers by ELISA. (**E**) PR8-specific total IgG antibody titers in sera from *Bcl6*^fl/fl^ (blue), *Bcl6*^fl/fl^*Cd4*^Cre^ (magenta), or *Ciita*^−/−^ (green) mice at 14 dpi with PR8. (**F**) PR8-specific IgG1, IgG2b, IgG2c, and IgG3 antibody titers at 14 dpi with PR8. LOD, limit of detection of the assay. Statistical significance was assessed by either two-tailed unpaired t-test or Welch’s t-test, based on the F test for unequal variance. \**P* < 0.05; \*\**P* < 0.01; \*\*\**P* < 0.001; \*\*\*\**P* < 0.0001. ns, not significant. Data are expressed as mean ± standard error of mean (SEM) log_10_ arbitrary units (AU). Each symbol represents an individual mouse. Data are aggregated from at least two independent experiments.

*Bcl6*^fl/fl^ mice produced high levels of spike (S)-specific IgG antibodies at 14 dpi (Fig. 1B). While *Bcl6*^fl/fl^*Cd4*^Cre^ mice had reduced levels of S-specific IgG, they still produced substantially more compared to *Ciita*^−/−^ mice. We next examined the requirement for Tfh cell help among different IgG subclasses. While S-specific IgG1 and IgG3 were completely Tfh-dependent, S-specific IgG2b and IgG2c were promoted by both Tfh and non-Tfh CD4^+^ T cells (Fig. 1C). Consistent with their divergent requirement for Tfh cell help, S-specific IgG2c was induced earlier than S-specific IgG1 (Fig. S1J). S-specific IgM was unaffected by the absence of Tfh cells (Fig. S1K). Together, these results suggest that both Tfh cells and non-Tfh CD4^+^ T cells promote the production of class-switched antibodies to SARS-CoV-2, though Tfh cells are uniquely able to induce certain subclasses.

We next asked whether these findings were generalizable to other models of respiratory viral infection. We infected mice with mouse-adapted influenza virus A/PR/8/34 H1N1 (PR8) and assessed antibody production at 14 dpi (Fig. 1D). Similar to SARS-CoV-2 infection, PR8 infection induced both Tfh-dependent and non-Tfh CD4^+^ T cell-dependent IgG antibodies (Fig. 1E). Again, PR8-specific IgG1 demonstrated a complete dependence on Tfh cell help, while PR8-specific IgG2b and IgG2c were promoted by both Tfh-dependent and -independent pathways (Fig. 1F). PR8-specific IgG3 and IgM were only partially dependent on Tfh and CD4^+^ T cell help (Fig. 1F and S1L). Thus, both Tfh and non-Tfh CD4^+^ T cells contribute to antibody production in two distinct models of respiratory viral infection.

### Th1 cells express Tfh effector molecules and interact with IgG2c^+^ B cells following viral infection

To determine which non-Tfh CD4^+^ T cell populations promote antibody production, we analyzed the medLN at 7 dpi with SARS-CoV-2. While total CD4^+^ T cell counts were unaffected in *Bcl6*^fl/fl^*Cd4*^Cre^ mice, activated CD44^+^CD4^+^ T cell counts were reduced, consistent with the loss of Tfh cells (Fig. S2, A and B). We classified PD-1^lo^CXCR5^lo^CD44^+^CD4^+^ T cells by their expression of PSGL-1 and Ly6C (Fig. S2C). These markers have previously been used in acute LCMV and influenza virus infection to distinguish terminally differentiated PSGL-1^hi^Ly6C^hi^ Th1 cells from a heterogeneous PSGL-1^hi^Ly6C^lo^ Th1 compartment containing memory precursors along with other Th1-related functional subsets (*38–40*). We therefore defined PSGL-1^hi^Ly6C^hi^ cells as terminally differentiated Th1 cells and PSGL-1^hi^Ly6C^lo^ cells as mixed Th1 cells. We also observed a BCL6-dependent PSGL-1^lo^Ly6C^lo^ population within the PD-1^lo^CXCR5^lo^ gate, which has previously been described as pre-Tfh cells (Fig. S2, D and E) (*41*). Mixed Th1 and terminally differentiated Th1 cells increased in relative frequency among activated CD4^+^ T cells in *Bcl6*^fl/fl^*Cd4*^Cre^ mice, consistent with the loss of Tfh and pre-Tfh cells; however, their absolute numbers were similar between *Bcl6*^fl/fl^ and *Bcl6*^fl/fl^*Cd4*^Cre^ mice, indicating that BCL6 deficiency did not lead to an aberrant increase in Th1 populations (Fig. S2, F to I).

We next evaluated whether these Th1 populations produce CD40L and IL-21, effector molecules usually ascribed to Tfh cells (*2*). CD40L and IL-21 act at multiple stages to support B cell activation, proliferation, differentiation, and antibody production (*2*). While Tfh cells produced the highest levels of CD40L, mixed Th1 cells also produced substantial levels of this effector molecule (Fig. 2, A and B). Tfh cells and mixed Th1 cells comprised the majority of CD40L-expressing CD4^+^ T cells in the medLN of *Bcl6*^fl/fl^ mice, while mixed Th1 cells became the main CD40L-expressing cells in *Bcl6*^fl/fl^*Cd4*^Cre^ mice (Fig. 2C). In *Bcl6*^fl/fl^*Cd4*^Cre^ mice, the frequency of CD40L^+^ cells among CD4^+^ T cells was decreased (Fig. S3A), likely owing to the loss of CD40L-expressing Tfh cells. However, mixed Th1 and terminally differentiated Th1 cells expressed higher levels of CD40L in *Bcl6*^fl/fl^*Cd4*^Cre^ mice compared to *Bcl6*^fl/fl^ mice (Fig. S3, B and C). In the absence of Tfh cells, Th1 subsets may have more opportunities to interact with antigen-presenting cells and experience T cell receptor signaling, which promotes CD40L expression (*42*). In *Bcl6*^fl/fl^ mice infected with PR8, mixed Th1 cells also expressed substantial levels of CD40L and constituted a large percentage of CD40L-expressing CD4^+^ T cells (Fig. S3, D and E). Therefore, mixed Th1 cells are a significant source of CD40L in the medLN of virally infected *Bcl6*^fl/fl^ and *Bcl6*^fl/fl^*Cd4*^Cre^ mice, suggesting that they may provide contact-dependent help to B cells both in the presence and in the absence of Tfh cells.

**Figure 2:**
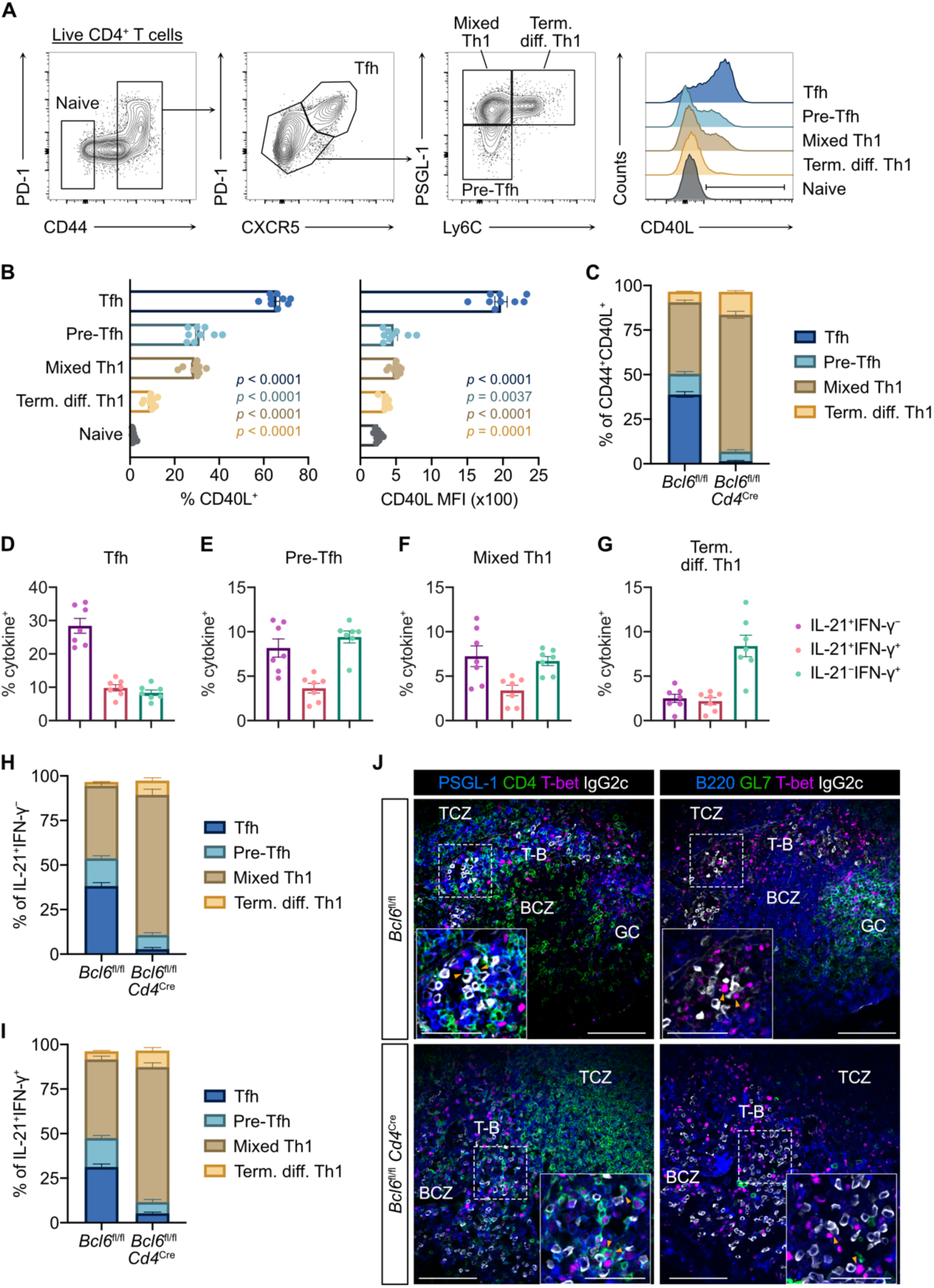
Th1 cells express Tfh effector molecules and interact with IgG2c^+^ B cells following viral infection. (**A** to **I**) Flow cytometric analysis of CD4^+^ T cells in mediastinal lymph nodes (medLN) from SARS-CoV-2-infected mice at 7 dpi. (**A**) Representative gating strategy for quantifying CD40L expression in distinct CD4^+^ T cell subsets. Pre-Tfh cells express a low amount of PD-1 and CXCR5 but are BCL6-dependent (*41*). Mixed Th1 cells classically include Th1 effectors and memory precursors. Terminally differentiated Th1 cells have been shown to express high levels of T-bet, IFN-γ, and GzmB (*38*). (**B**) Frequency (left) and median fluorescence intensity (MFI) (right) of CD40L expression within CD4^+^ T cell subsets from control *Bcl6*^fl/fl^ mice. Statistical significance was assessed by two-tailed unpaired Welch’s t-test. *P* values for each subset relative to naive CD4^+^ T cells are color-coded. (**C**) Relative proportions of CD4^+^ T cell subsets among CD44^+^CD40L^+^ cells from *Bcl6*^fl/fl^ or *Bcl6*^fl/fl^*Cd4*^Cre^ mice. (**D** to **G**) Frequency of IL-21^+^IFN-γ^−^ (purple), IL-21^+^IFN-γ^+^ (red), and IL-21^−^IFN-γ^+^ (teal) expression by intracellular cytokine staining in Tfh cells (D), pre-Tfh cells (E), mixed Th1 cells (F), and terminally differentiated Th1 cells (G) cells from control *Bcl6*^fl/fl^ mice. (**H** and **I**) Relative proportions of CD4^+^ T cell subsets among IL-21^+^IFN-γ^−^ (H) and IL-21^+^IFN-γ^+^ (I) cells from *Bcl6*^fl/fl^ or *Bcl6*^fl/fl^*Cd4*^Cre^ mice. Data are expressed as mean ± SEM. Each symbol represents an individual mouse. Data are aggregated from at least two independent experiments. (**J**) Immunofluorescence of serial sections of medLN from *Bcl6*^fl/fl^ (top row) or *Bcl6*^fl/fl^*Cd4*^Cre^ (bottom row) mice infected with PR8 at 14 dpi. Left column: PSGL-1 (blue), CD4 (green), T-bet (magenta), and IgG2c (white). Right column: B220 (blue), GL7 (green), T-bet (magenta), and IgG2c (white). Dashed line demarcates region shown in inset. Orange arrowheads denote Th1 cells interacting with IgG2c^+^ B cells. BCZ, B cell zone. TCZ, T cell zone. T-B, T cell–B cell border. GC, germinal center. Representative images from 3 mice; scale bars, 100 µm and 50 µm (inset).

In SARS-CoV-2 infection, Tfh cells also produced the highest levels of IL-21 (Fig. 2D and S3F). A portion of Tfh cells produced IL-21 together with IFN-γ, which is important for IgG2c class switching (*43, 44*). Pre-Tfh cells and mixed Th1 cells consisted of IL-21 single producers, IFN-γ single producers, and IL-21/IFN-γ double producers (Fig. 2, E and F). Terminally differentiated Th1 cells were predominantly IFN-γ single producers (Fig. 2G), suggesting that they likely migrate to peripheral tissues to exert their effector function (*38*). As with CD40L expression, Tfh cells and mixed Th1 cells comprised the majority of IL-21 single-producing and IL-21/IFN-γ double-producing CD4^+^ T cells in *Bcl6*^fl/fl^ mice (Fig. 2, H and I). In *Bcl6*^fl/fl^*Cd4*^Cre^ mice, the frequency of IL-21 single producers was decreased, potentially due to the loss of Tfh cells, and mixed Th1 cells became the dominant cytokine-producing population (Fig. S3G and Fig. 2, H and I). Mixed Th1 and terminally differentiated Th1 cells displayed similar cytokine profiles between *Bcl6*^fl/fl^ and *Bcl6*^fl/fl^*Cd4*^Cre^ mice (Fig. S3, H and I).

As mixed Th1 cells express IL-21 in addition to CD40L, they could support an alternative pathway of antibody production to that driven by Tfh cells. We therefore assessed whether mixed Th1 cells are sub-anatomically positioned to provide help to B cells. PSGL-1 mediates chemotaxis to CCL21 and CCL19, therefore helping naïve CD4^+^ T cells home to secondary lymphoid organs (*45*). During their differentiation, Tfh cells downregulate PSGL-1 and upregulate CXCR5 to enable their migration into B cell follicles (*41, 46*). However, as mixed Th1 cells continue to express PSGL-1 and do not upregulate CXCR5, it is unclear whether they can migrate to sites of B cell help. Immunofluorescence of medLN following PR8 infection showed that Th1 cells (PSGL-1^+^CD4^+^T-bet^+^) co-localized with IgG2c^+^ B cells at the T cell-B cell (T-B) border (Fig. 2J). This was true in both *Bcl6*^fl/fl^ and *Bcl6*^fl/fl^*Cd4*^Cre^ mice, suggesting that Th1 cells promote antibody production in parallel with Tfh cells as well as in their absence. Taken together, we found that a subset of mixed Th1 cells expressed CD40L and IL-21 and were positioned at the T-B border to help B cells during viral infection. To distinguish this population from the Th1 cells that function in peripheral tissues (e.g., the lung), we call these cells “LN-Th1” cells.

### Durable, high-affinity antibodies to SARS-CoV-2 are made in the absence of Tfh cells

We next measured the affinity of the antibodies generated during SARS-CoV-2 and PR8 infection. As GCs are the conventional site of somatic hypermutation and affinity maturation (*2*), we expected that *Bcl6*^fl/fl^ mice would generate high-affinity antibodies through Tfh/GC-dependent processes while *Bcl6*^fl/fl^*Cd4*^Cre^ mice would generate low-affinity antibodies through LN-Th1-driven responses. *Bcl6*^fl/fl^ mice produced high-affinity IgG antibodies to S as well as the spike receptor-binding domain (RBD) (Fig. 3A), a major target of neutralizing antibodies (*47*). In contrast, S- and RBD-specific IgG antibodies from *Ciita*^−/−^ mice displayed minimal affinity. However, we discovered that antibodies from *Bcl6*^fl/fl^*Cd4*^Cre^ mice still demonstrated substantial affinity toward S and RBD, suggesting that LN-Th1 cells could promote high-affinity antibody production.

**Figure 3:**
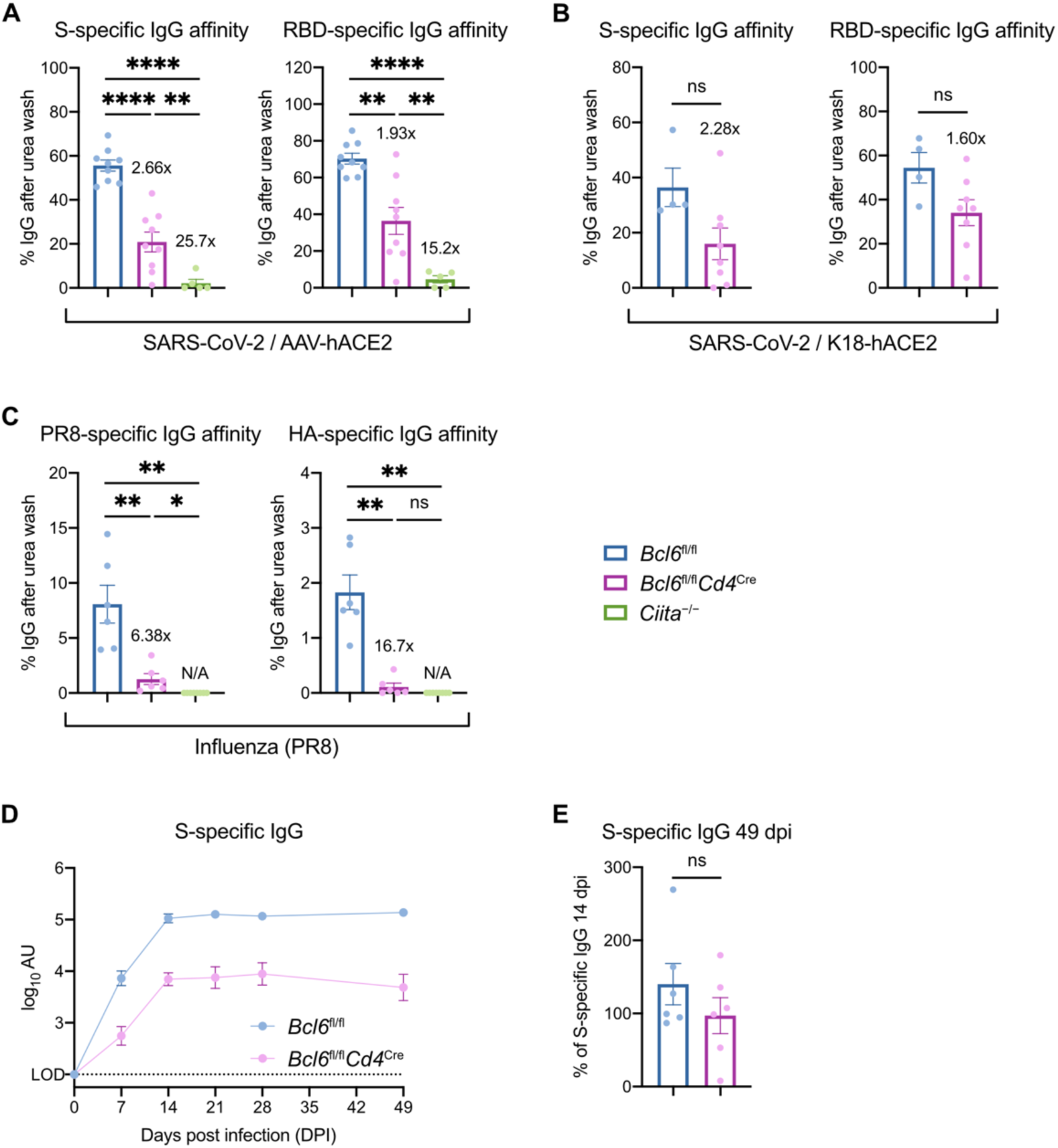
Durable, high-affinity antibodies to SARS-CoV-2 are made in the absence of Tfh cells. (**A**) Affinity index of serum IgG antibodies against SARS-CoV-2 S (left) or RBD (right), calculated as relative ELISA signal after urea wash. Sera are from *Bcl6*^fl/fl^ (blue), *Bcl6*^fl/fl^*Cd4*^Cre^ (magenta), or *Ciita*^−/−^ (green) mice infected with SARS-CoV-2 at 14 dpi. Fold changes relative to *Bcl6*^fl/fl^ mice are annotated. (**B**) Affinity index of serum IgG antibodies against SARS-CoV-2 S (left) or RBD (right). Sera are from K18-hACE2 *Bcl6*^fl/fl^ (blue) or K18-hACE2 *Bcl6*^fl/fl^*Cd4*^Cre^ (magenta) mice infected with SARS-CoV-2 at 14 dpi. Fold changes relative to K18-hACE2 *Bcl6*^fl/fl^ mice are annotated. (**C**) Affinity index of serum IgG antibodies against PR8 (left) or HA (right), calculated as relative ELISA signal after urea wash. Sera are from *Bcl6*^fl/fl^ (blue), *Bcl6*^fl/fl^*Cd4*^Cre^ (magenta), or *Ciita*^−/−^ (green) mice infected with PR8 at 14 dpi. Fold changes relative to *Bcl6*^fl/fl^ mice are annotated. (**D**) Longitudinal dynamics of S-specific IgG antibodies up to 49 dpi, in sera from *Bcl6*^fl/fl^ or *Bcl6*^fl/fl^*Cd4*^Cre^ mice infected with SARS-CoV-2. (**E**) Relative serum titers of S-specific IgG at 49 dpi, compared to S-specific IgG at 14 dpi. Statistical significance was assessed by either two-tailed unpaired t-test or Welch’s t-test, based on the F test for unequal variance. \**P* < 0.05; \*\**P* < 0.01; \*\*\*\**P* < 0.0001. ns, not significant. Data are expressed as mean ± SEM. Each symbol in (A to C and E) represents an individual mouse. Each symbol in (D) represents the mean of six mice. Data are aggregated from at least two independent experiments.

To determine whether this was also true in a SARS-CoV-2 infection model that does not require AAV pre-transduction, we crossed *Bcl6*^fl/fl^ and *Bcl6*^fl/fl^*Cd4*^Cre^ mice to K18-hACE2 transgenic mice, which express hACE2 in epithelial cells (*48*). Both K18-hACE2 *Bcl6*^fl/fl^ and *Bcl6*^fl/fl^*Cd4*^Cre^ mice produced high-affinity antibodies to S and RBD (Fig. 3B), indicating that non-Tfh cells can support high-affinity antibody production in two separate models of SARS-CoV-2 infection. However, this was not the case with PR8 infection, as *Bcl6*^fl/fl^*Cd4*^Cre^ mice produced IgG antibodies of minimal affinity to both PR8 and PR8 surface glycoprotein hemagglutinin (HA) (Fig. 3C). Therefore, the ability of non-Tfh cells, likely LN-Th1 cells, to promote high-affinity antibodies may depend on the nature of the viral infection and the antigenic target.

We also evaluated the durability of S-specific IgG antibodies produced by *Bcl6*^fl/fl^ and *Bcl6*^fl/fl^*Cd4*^Cre^ mice following SARS-CoV-2 infection. Previous studies of viral infection have shown that antibody titers in mice with impaired Tfh cells are especially reduced at later timepoints (*18, 19, 25*). In SARS-CoV-2 infection, S-specific IgG levels peaked at 14 dpi in *Bcl6*^fl/fl^ mice and were stable through 49 dpi (Fig. 3D). In *Bcl6*^fl/fl^*Cd4*^Cre^ mice, S-specific IgG antibodies were reduced tenfold at all timepoints measured yet remained stable between 14 dpi and 49 dpi (Fig. 3E). Together, these results indicate that Tfh-independent antibodies to SARS-CoV-2 are both high-affinity and durable – two important qualities usually attributed to Tfh-dependent responses.

### Tfh cells focus the antibody repertoire but are dispensable for broad coverage of SARS-CoV-2 epitopes

We next characterized the antibody epitope repertoire of Tfh-versus LN-Th1-driven responses to SARS-CoV-2. Sera from AAV-hACE2 *Bcl6*^fl/fl^ and *Bcl6*^fl/fl^*Cd4*^Cre^ mice at 14 dpi were profiled using a bacterial display library of 2410 linear peptides tiling the entire SARS-CoV-2 proteome (Fig. 4A). We first compared the diversity of antibody epitope reactivity, calculating the Shannon entropy, Simpson’s diversity index, and the repertoire focusing index within each sample (Methods). We observed that antibody diversity assessed by Shannon entropy and Simpson’s diversity index was significantly decreased in *Bcl6*^fl/fl^ mice compared to *Bcl6*^fl/fl^*Cd4*^Cre^ mice, while the degree of repertoire focusing was increased (Fig. 4B). These findings were robust to variations in read counts (Fig. S4A). These results suggest that Tfh cells help focus the antibody response to particular viral epitopes while LN-Th1 cells promote antibodies to a wider array of targets.

**Figure 4:**
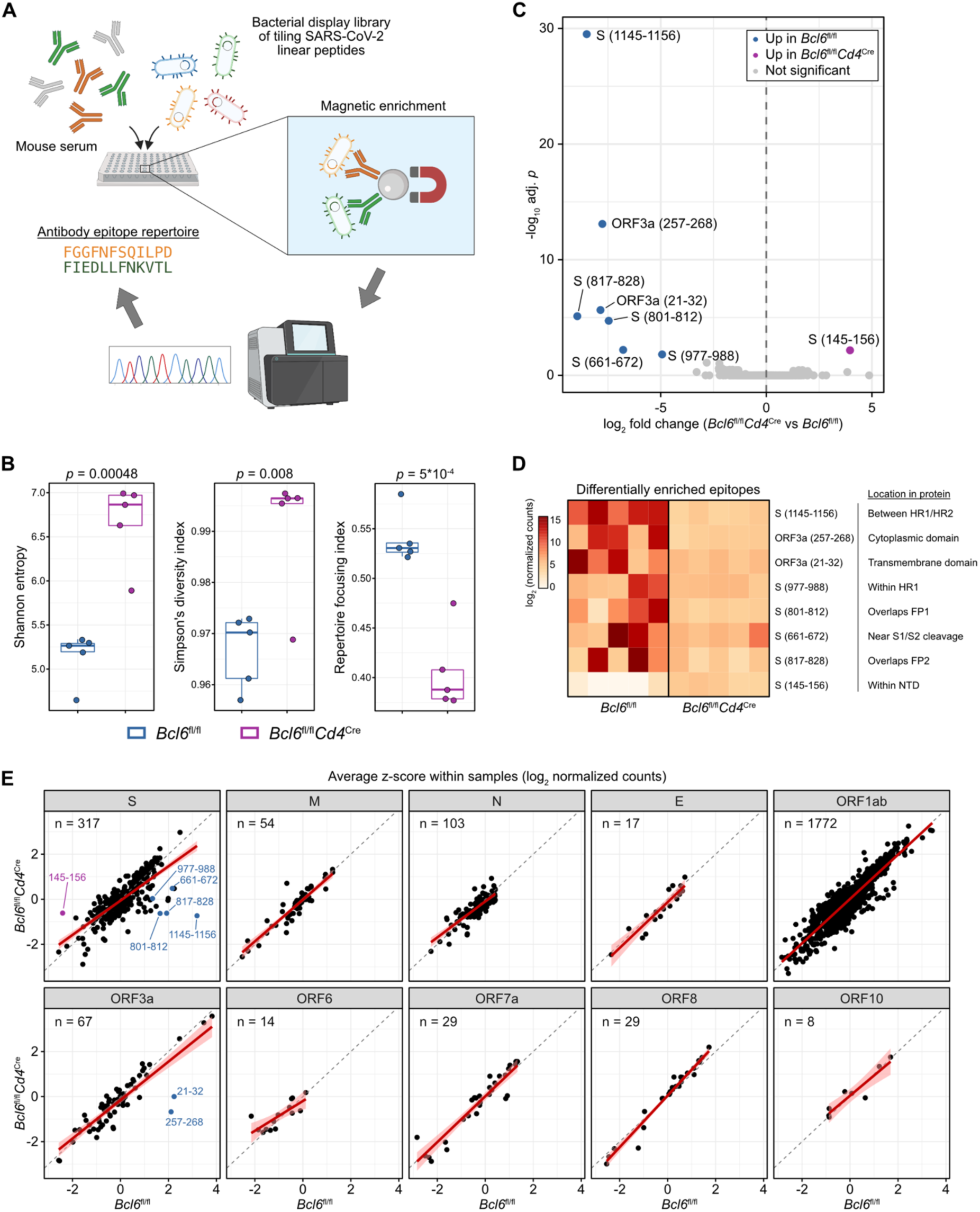
Tfh cells focus the antibody repertoire but are dispensable for broad coverage of SARS-CoV-2 epitopes. (**A**) Sera from infected *Bcl6*^fl/fl^ or *Bcl6*^fl/fl^*Cd4*^Cre^ mice (*n* = 5 from two independent experiments) were assayed for reactivity against a bacterial display library of overlapping linear epitopes tiling the SARS-CoV-2 proteome. (**B**) Tukey boxplots detailing Shannon entropy (left), Simpson’s diversity index (middle), and repertoire focusing index (right) in *Bcl6*^fl/fl^ or *Bcl6*^fl/fl^*Cd4*^Cre^ mice. Statistical significance was assessed by two-tailed unpaired Welch’s t-test. Each symbol represents an individual mouse. (**C**) Volcano plot of differentially enriched linear epitopes in *Bcl6*^fl/fl^ vs *Bcl6*^fl/fl^*Cd4*^Cre^ mice. Epitopes that are significantly enriched in *Bcl6*^fl/fl^ sera are colored in blue, while epitopes that are enriched in *Bcl6*^fl/fl^*Cd4*^Cre^ sera are colored in magenta. Point labels describe the amino acids within the indicated SARS-CoV-2 protein from which the epitope was derived. Statistical significance was assessed by two-tailed DESeq2 Wald test, with Benjamini-Hochberg multiple hypothesis correction (adjusted *p* < 0.05). (**D**) Heatmap of differentially enriched linear epitopes in *Bcl6*^fl/fl^ vs *Bcl6*^fl/fl^*Cd4*^Cre^ mice, from (C). Data are expressed as log_2_-transformed normalized counts. The position of each epitope in relation to known protein domains is annotated. (**E**) Scatter plots of relative enrichment scores for linear epitopes derived from each SARS-CoV-2 protein, comparing *Bcl6*^fl/fl^ vs *Bcl6*^fl/fl^*Cd4*^Cre^ sera. Data are expressed as average *z-*scores (log_2_ normalized counts were scaled to *z*-scores within each sample, then averaged across *Bcl6*^fl/fl^ or *Bcl6*^fl/fl^*Cd4*^Cre^ mice). The gray dashed line demarcates equivalence between *Bcl6*^fl/fl^ and *Bcl6*^fl/fl^*Cd4*^Cre^, while the red line denotes the linear regression model with 95% confidence intervals shaded in pink. Epitopes that were identified as differentially enriched are annotated as in (C). The number of epitopes derived from each SARS-CoV-2 protein is annotated in the top left.

Given these changes in antibody diversity, we next explored whether there were differences in antibody reactivity at the level of individual linear epitopes. After normalizing for read count variations using the median of ratios method (Fig. S4B), we identified epitopes that were comparatively enriched or depleted in *Bcl6*^fl/fl^ versus *Bcl6*^fl/fl^*Cd4*^Cre^ mice (Fig. 4C). We found that seven epitopes were enriched in *Bcl6*^fl/fl^ mice, while one epitope was depleted (Fig. 4D). Five of the seven enriched epitopes were derived from S: aa661-672 (proximal to S1/S2 cleavage site), aa801-812 (fusion peptide [fp] 1), aa817-828 (FP2), aa977-988 (heptad repeat [HR] 1), and aa1145-1156 (between HR1/HR2). On the other hand, aa145-156 (N-terminal domain) from S was comparatively depleted in *Bcl6*^fl/fl^ mice.

To further investigate alterations in epitope reactivity, we converted the normalized counts to z-scores on a sample-by-sample basis, such that the z-scores would denote the relative rank of a specific epitope within a particular sample (Fig. S4C). Consistent with our prior analyses, the average z-scores in *Bcl6*^fl/fl^ versus *Bcl6*^fl/fl^*Cd4*^Cre^ mice were similar across most SARS-CoV-2 proteins, with the exception of regions within the S and ORF3a proteins (Fig. 4E). In particular, the regression lines for non-S proteins all closely followed the line of identity, indicating that the relative ranks of epitopes from non-S proteins were largely similar in the presence or absence of Tfh cells. These analyses therefore indicate that, while LN-Th1 cells can promote antibody production against most SARS-CoV-2 epitopes, Tfh cells focus the antibody response against certain S-derived epitopes.

### RBD-specific antibodies are generated in the absence of Tfh cells

Analyzing antibody epitope reactivity along the length of S, we observed that the majority of epitopes enriched in *Bcl6*^fl/fl^ mice (17/21 epitopes with differential average z-score > 1) were found in the S2 domain (aa686-1273; Fisher’s exact test *p* = 0.0012) (Fig. 5A), which mediates fusion of viral and target cell membranes (*49*). These included most of the aforementioned S-derived epitopes that were significantly enriched (Fig. 4C), as well as contiguous epitopes whose enrichment did not reach statistical significance in the epitope-level analysis. For example, several epitopes in the fusion peptide were enriched adjacent to the significantly enriched epitopes aa801-812 and aa817-828 (Fig. S4D). Many of these epitopes are highly conserved across human coronaviruses (hCoVs) as well as the emerging variants of concern (Fig. 5A). Interestingly, the enriched epitopes spanning FP1/FP2 and preceding HR2 have also been identified in numerous studies profiling the antibody epitope repertoire of COVID-19 patients (*50–60*). Given their immunodominance and conservation across hCoVs, these epitopes have been proposed as targets for a pan-coronavirus vaccine, though it remains unclear whether antibodies directed against these regions are actually able to neutralize virus (*51, 52, 55, 57, 58, 61*).

**Figure 5:**
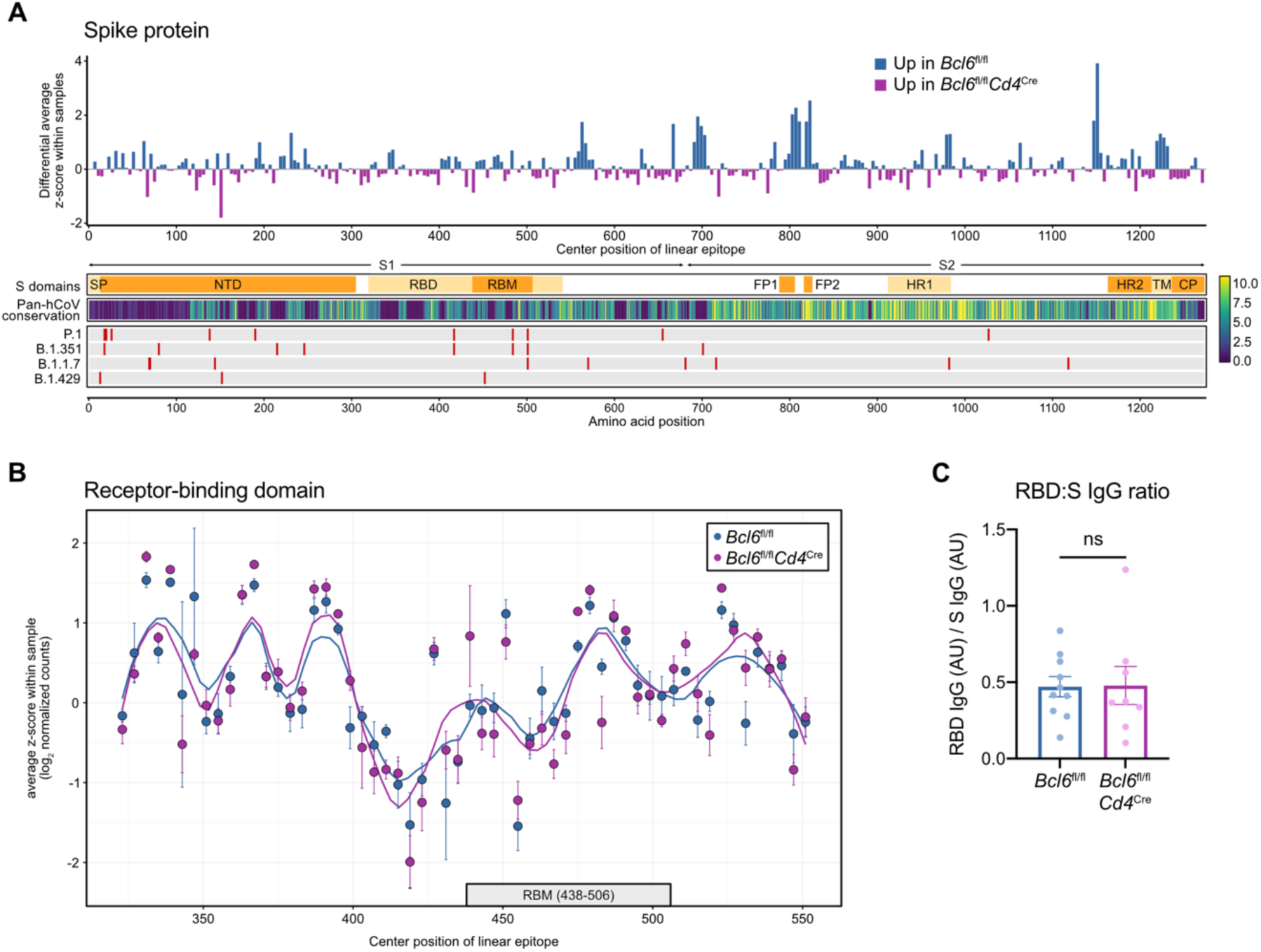
RBD-specific antibodies are generated in the absence of Tfh cells. (**A**) Top: differential enrichment scores for spike (S)-derived linear epitopes, such that epitopes that are comparatively enriched in *Bcl6*^fl/fl^ mice are represented by positive values (blue), whereas epitopes that are relatively enriched in *Bcl6*^fl/fl^*Cd4*^Cre^ mice are indicated by negative values (magenta). Linear epitopes are each indicated by their center position within the S protein. Bottom: annotation tracks detailing S protein domains, pan-human coronavirus (hCoV) conservation scores, and mutations found in select SARS-CoV-2 variants of concern. SP, signal peptide. NTD, N-terminal domain. RBD, receptor-binding domain. RBM, receptor-binding motif. FP1, fusion peptide 1. FP2, fusion peptide 2. HR1, heptad repeat 1. HR2, heptad repeat 2. TM, transmembrane domain. CP, cytoplasmic domain. (**B**) Relative enrichment scores for linear epitopes derived from RBD, comparing *Bcl6*^fl/fl^ vs *Bcl6*^fl/fl^*Cd4*^Cre^ sera. Data are expressed as average *z-*scores in *Bcl6*^fl/fl^ or *Bcl6*^fl/fl^*Cd4*^Cre^ mice, with SEM error bars and loess regression lines. The RBM is annotated below with a gray box. (**C**) Ratio of RBD-specific to S-specific IgG antibodies in sera from *Bcl6*^fl/fl^ or *Bcl6*^fl/fl^*Cd4*^Cre^ mice at 14 dpi with SARS-CoV-2. Statistical significance was assessed by two-tailed unpaired t-test. ns, not significant. Data are expressed as mean ± SEM. Each symbol represents an individual mouse. Data are aggregated from three independent experiments.

In contrast, *Bcl6*^fl/fl^ and *Bcl6*^fl/fl^*Cd4*^Cre^ mice demonstrated similar antibody reactivity to most epitopes within the RBD (Fig. 5B), the target of most neutralizing antibodies (*62*). However, as most antibodies to RBD likely recognize conformational epitopes (*56, 57*), we also measured RBD-specific antibodies by ELISA using full-length RBD. RBD-specific IgG titers normalized by total S-specific IgG were similar between *Bcl6*^fl/fl^ and *Bcl6*^fl/fl^*Cd4*^Cre^ mice (Fig. 5C). Thus, while Tfh cells focus the antibody response against immunodominant S2 epitopes, LN-Th1 cells still promote antibodies against the primary target of neutralization, RBD.

### Tfh-dependent and -independent antibodies demonstrate similar neutralization potency against homologous SARS-CoV-2 as well as the B.1.351 variant of concern

We next evaluated the function of antibodies from *Bcl6*^fl/fl^ and *Bcl6*^fl/fl^*Cd4*^Cre^ mice following SARS-CoV-2 infection. While we had observed that Tfh-independent antibody responses lacked IgG1/IgG3 subclasses (Fig. 1C) and S2 epitope focusing (Fig. 5A), these antibodies were still high-affinity (Fig. 3, A and B) and could target the RBD (Fig. 5C). We therefore expected that Tfh-independent, LN-Th1-driven antibodies would demonstrate similar neutralizing function against homologous SARS-CoV-2 (USA-WA1/2020) as those generated with Tfh cell help. Using vesicular stomatitis virus (VSV) pseudotyped with USA-WA1/2020 S protein, we measured the neutralization titer (the reciprocal serum dilution achieving 50% neutralization of pseudovirus infection, NT50) of sera from *Bcl6*^fl/fl^ and *Bcl6*^fl/fl^*Cd4*^Cre^ mice. *Bcl6*^fl/fl^ sera exhibited increased NT50 (Fig. 6A), which was expected given their higher levels of S-specific IgG antibodies (Fig. 1B). However, by normalizing NT50 to S-specific IgG levels in each sample, we observed that the neutralization potency indices of *Bcl6*^fl/fl^ and *Bcl6*^fl/fl^*Cd4*^Cre^ sera were similar and actually trended higher for *Bcl6*^fl/fl^*Cd4*^Cre^ sera (Fig. 6A).

**Figure 6:**
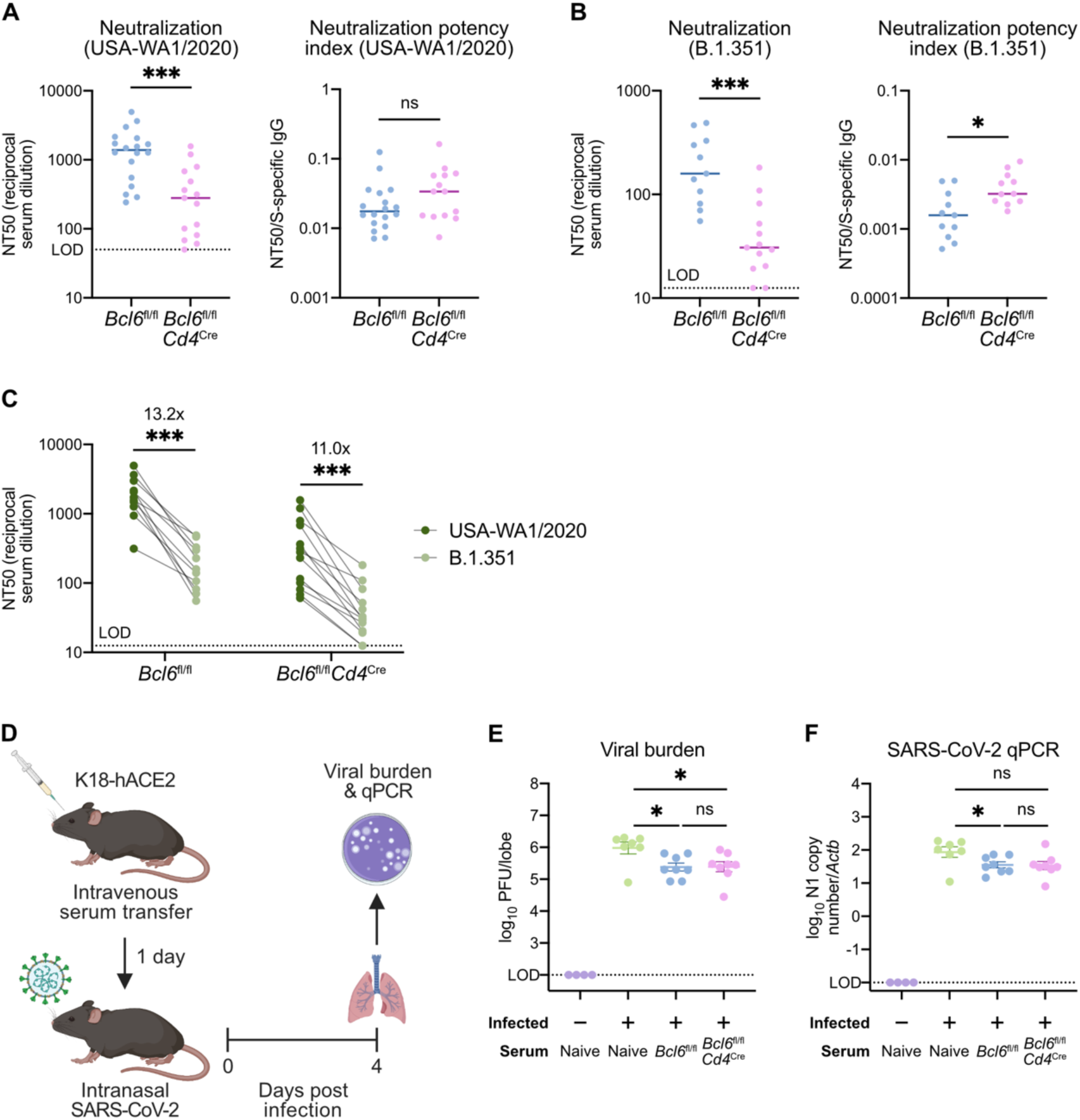
Tfh-dependent and -independent antibodies demonstrate similar neutralization potency against homologous SARS-CoV-2 as well as the B.1.351 variant of concern. (**A**) Left: neutralizing titers (inverse of half maximal inhibitory concentration; NT50) of sera from *Bcl6*^fl/fl^ (blue) or *Bcl6*^fl/fl^*Cd4*^Cre^ (magenta) mice against VSV particles pseudotyped with homologous S protein (USA-WA1/2020, the same isolate used to infect the mice). Right: neutralization potency indices (NT50 divided by S-specific total IgG) of sera from *Bcl6*^fl/fl^ or *Bcl6*^fl/fl^*Cd4*^Cre^ mice against USA-WA1/2020 pseudovirus. (**B**) Left: neutralizing titers of sera from *Bcl6*^fl/fl^ (blue) or *Bcl6*^fl/fl^*Cd4*^Cre^ (magenta) mice against VSV particles pseudotyped with variant S protein (B.1.351). Right: neutralization potency indices of sera from *Bcl6*^fl/fl^ or *Bcl6*^fl/fl^*Cd4*^Cre^ mice against B.1.351 pseudovirus. (**C**) Matched comparison of neutralizing titers against homologous (USA-WA1/2020) vs variant (B.1.351) pseudovirus. Fold changes are indicated above each group. (**D**) Schematic of experimental design for assessing *in vivo* neutralization potency of sera. (**E** and **F**) Viral burden in lungs from K18-hACE2 mice infected with SARS-CoV-2 at 4 dpi or left uninfected. Mice were treated with serum from naive, infected *Bcl6*^fl/fl^, or infected *Bcl6*^fl/fl^*Cd4*^Cre^ mice. Data are expressed as log_10_ plaque forming units (PFU) per lung lobe by plaque assay (E) or log_10_ N1 gene copy number by qPCR, normalized to *Actb* (F). Statistical significance was assessed by two-tailed Mann–Whitney test (A and B), two-tailed Wilcoxon signed-rank test (C), or two-tailed unpaired t-test (E and F). \**P* < 0.05; \*\*\**P* < 0.001. ns, not significant. Data are expressed as the median for (**A** and **B**) and expressed as mean ± SEM for (**E** and **F**). Each symbol represents an individual mouse. Data are aggregated from at least two independent experiments.

We next tested the same sera against VSV pseudotyped with S protein from the B.1.351 variant of concern. Multiple studies have shown that B.1.351 S mutations, particularly those in the RBD, disrupt binding by neutralizing antibodies and facilitate immune escape (*63–68*). We therefore hypothesized that increased focusing of Tfh-dependent antibodies against conserved S2 epitopes would enable *Bcl6*^fl/fl^ sera to better neutralize B.1.351 pseudovirus than *Bcl6*^fl/fl^*Cd4*^Cre^ sera. While *Bcl6*^fl/fl^ sera exhibited greater NT50 than *Bcl6*^fl/fl^*Cd4*^Cre^ sera, we found that the neutralization potency index of *Bcl6*^fl/fl^*Cd4*^Cre^ sera was higher than that of *Bcl6*^fl/fl^ sera (Fig. 6B). Nevertheless, both *Bcl6*^fl/fl^ and *Bcl6*^fl/fl^*Cd4*^Cre^ sera demonstrated an approximately 10-fold reduction in NT50 against the B.1.351 variant compared to USA-WA1/2020 (Fig. 6C), consistent with previous studies (*64, 67, 68*). Our findings therefore indicate that Tfh-independent antibodies exhibit similar, if not increased, neutralization potency as Tfh-dependent antibodies *in vitro* and that S2 epitope focusing does not improve neutralization activity against the B.1.351 variant of concern.

Finally, we assessed the function of *Bcl6*^fl/fl^ and *Bcl6*^fl/fl^*Cd4*^Cre^ sera against SARS-CoV-2 *in vivo*. To this end, we transferred S-specific IgG titer-matched sera from *Bcl6*^fl/fl^ or *Bcl6*^fl/fl^*Cd4*^Cre^ mice infected with SARS-CoV-2 (USA-WA1/2020) into naïve K18-hACE2 mice (Fig. 6D). One day later, we infected the K18-hACE2 recipients with homologous SARS-CoV-2 and then measured viral burden in the lungs at 4 dpi. *Bcl6*^fl/fl^ and *Bcl6*^fl/fl^*Cd4*^Cre^ sera similarly reduced lung viral burden (Fig. 6, E and F). Taken together, these results demonstrate that Tfh-independent antibodies efficiently neutralize both homologous SARS-CoV-2 and the B.1.351 variant of concern and are functional both *in vitro* and *in vivo*.

## Discussion

While protective antibodies are usually generated through Tfh/GC-dependent pathways, it is unclear what happens to the antibody response when these structures are disrupted by virus-induced inflammation. We found that certain class-switched antibodies were reduced but still present in Tfh-deficient mice during both SARS-CoV-2 and influenza virus infection. Tfh-independent antibodies to SARS-CoV-2, likely driven by LN-Th1 cells, were still high-affinity and durable while also demonstrating more diverse epitope reactivity compared to Tfh-dependent antibodies. Importantly, LN-Th1-driven antibody responses neutralized both homologous SARS-CoV-2 (USA-WA1/2020) and the B.1.351 variant of concern and were functional *in vitro* and *in vivo* (Fig. 7).

**Figure 7:**
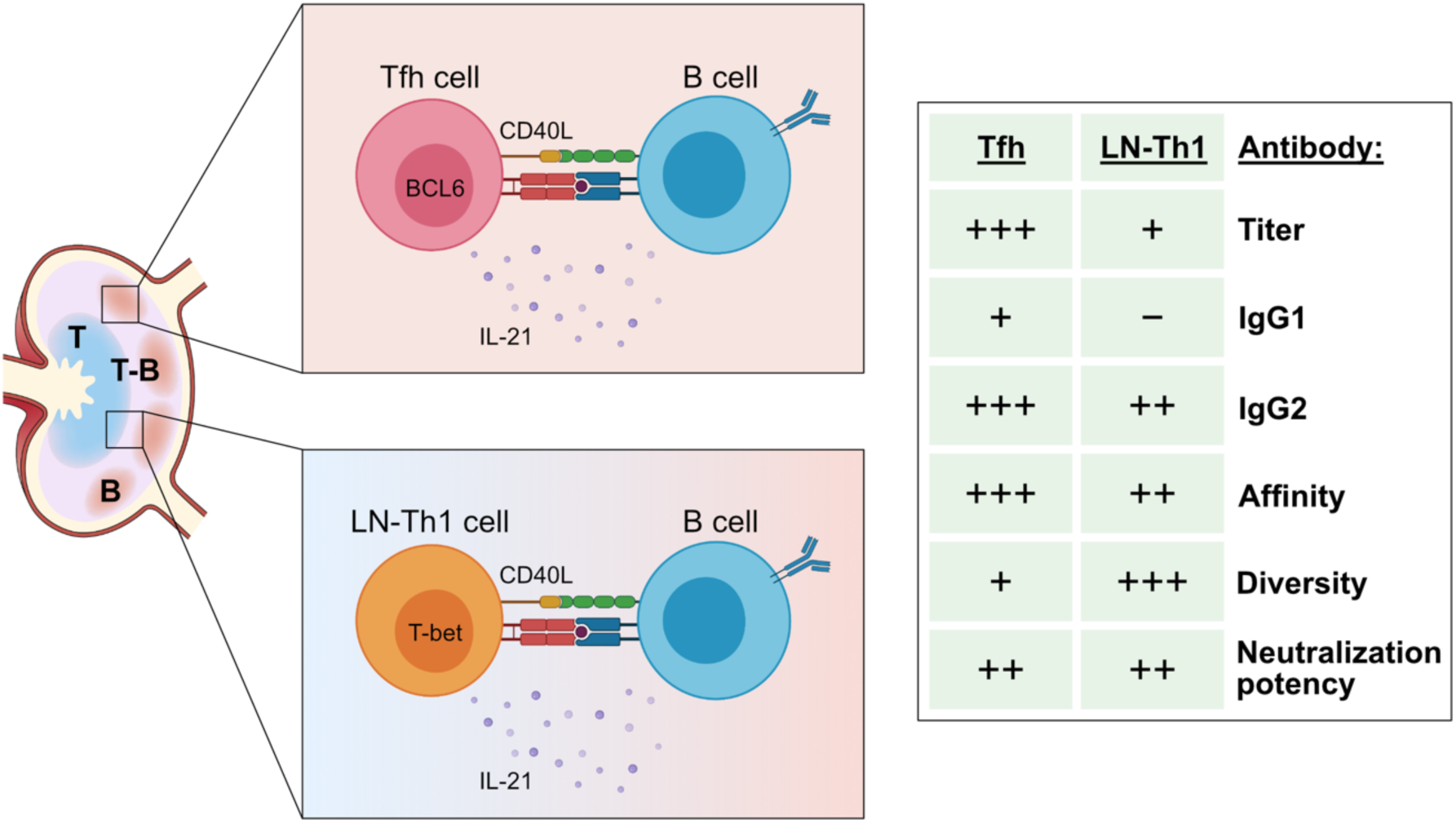
Tfh cells and LN-Th1 cells mediate complementary pathways of neutralizing antibody production. Tfh cells are located in follicles and germinal centers of lymph nodes where they provide cues for B cell class switching and selection, in part through CD40L and IL-21 signals. Similarly, lymph node (LN)-Th1 cells express CD40L and IL-21 but are primarily located at the T cell-B cell border of the lymph node. While LN-Th1-driven antibody responses are lower in titer and lack IgG1 class-switching, they generate IgG2 antibodies of high affinity, more diverse epitope reactivity, and similar neutralization potency. T, T cell zone. T-B, T cell-B cell border. B, B cell zone.

Though class-switched antibodies could be produced in the absence of Tfh cells, we observed that certain subclasses demonstrated a complete dependence on Tfh cell help. For example, IgG1 production was completely abrogated in Tfh-deficient mice infected with SARS-CoV-2 or influenza virus. This is consistent with previous studies analyzing antibody production by Tfh-deficient mice in the setting of vaccination or Zika virus infection (*25, 27*). In contrast, IgG2c was still made without Tfh cells but generally required CD4^+^ T cell help. This divergent requirement for Tfh cell help may be explained by the different sources of class-switch factors IL-4 and IFN-γ in the medLN. During viral infection, Tfh cells are the primary CD4^+^ T cell population that produces IL-4, a critical switch factor for IgG1 (*43, 69*). On the other hand, we found that, in addition to Tfh cells, Th1 cells can make IFN-γ along with IL-21 to promote IgG2c antibody production during SARS-CoV-2 infection. IFN-γ and IL-21 production by Th1 cells has also been observed in the setting of influenza vaccination (*27*). IgG1 and IgG2c exhibit distinct Fc effector functions as IgG1 preferentially binds the inhibitory Fc receptor (FcR) FcγRIIB, while IgG2c preferentially binds activating FcRs (*70*). Though IgG1 and IgG2 antibodies with the same Fab similarly neutralize SARS-CoV-2 *in vitro*, IgG2 Fc effector function is critical for viral clearance *in vivo* (*71, 72*). Thus, we speculate that the ability of LN-Th1 cells to promote IgG2c antibodies ensures that sufficient levels of antibodies with activating Fc effector function are produced, both in the presence and absence of Tfh cells.

We also demonstrated that LN-Th1 cells interact with IgG2c^+^ B cells at the T-B border. This is a plausible location for LN-Th1-driven antibody responses, as previous studies have shown that antigen-specific B cell proliferation occurs at the T-B border in addition to the GC (*73*). PSGL-1 expression on naïve CD4^+^ T cells is required for chemotaxis to CCL21 and CCL19, thus mediating homing to secondary lymphoid organs and ultimately the T cell zone (*41, 45*). Once activated, T cells upregulate several glycosyltransferases that modify PSGL-1 such that it can interact with P-selectin and enable trafficking to sites of inflammation (*74, 75*). These carbohydrate modifications of PSGL-1 also interfere with its ability to interact with CCL21 (*45*), providing a possible mechanism for how PSGL-1^hi^ LN-Th1 cells migrate from the T cell zone to the T-B border. Upregulation of chemokine receptors such as CXCR3 may also support LN-Th1 migration, as CXCR3 ligands CXCL9 and CXCL10 have been shown to regulate intranodal positioning and optimal differentiation of Th1 cells (*76, 77*). However, in contrast to terminally differentiated Th1 cells that then emigrate from the LN to coordinate cellular responses in peripheral tissues, LN-Th1 cells stay in the LN to provide B cell help and promote antibody production.

A surprising finding from our work was that SARS-CoV-2, but not influenza virus, induced high-affinity Tfh-independent antibodies. However, CD4^+^ T cell-independent antibodies demonstrated minimal affinity, indicating that either Tfh or LN-Th1 cells are required for high-affinity responses. While affinity maturation conventionally occurs in the GC through the iterative process of somatic hypermutation and competition for Tfh cell help (*2*), there are several possible mechanisms by which LN-Th1-driven antibody responses could achieve high affinity. Somatic hypermutation has also been shown to occur at extrafollicular sites during both infection and chronic autoimmunity (*78–80*), and LN-Th1 cells could therefore help select B cells that acquire affinity-increasing BCR mutations. Alternatively, multiple studies have identified potent, neutralizing antibodies with minimal somatic hypermutation from COVID-19 patients, suggesting that the naïve repertoire in humans already harbors protective antibodies that do not require additional affinity maturation (*81–89*). Mice may also possess near-germline antibodies with high affinity to SARS-CoV-2, and therefore the role of LN-Th1 cells in this case would be to select and expand the B cells expressing these sequences.

Epitope profiling revealed that Tfh cells focus the antibody repertoire against S2-derived epitopes that are highly conserved across human coronaviruses as well as the emerging variants of concern. These same epitopes spanning FP1/FP2 and preceding HR2 have been repeatedly identified in studies profiling the antibody repertoire of COVID-19 patients, suggesting that the immunodominance of these epitopes in humans is mediated by Tfh cells. However, it is less clear whether these S2-reactive antibodies are actually neutralizing. Analyses of S2-reactive antibodies from patient sera demonstrate that these antibodies have weak or absent neutralizing function (*51, 52, 55, 89*). Additionally, vaccination studies in rabbits and mice with S2 or S2-derived epitopes have reached conflicting conclusions about the ability of the resultant sera to neutralize SARS-CoV-2 (*57, 58, 90*). SARS-CoV-2 S2-cross-reactive antibodies have also been identified in pre-pandemic serum samples, likely induced by exposure to seasonal coronaviruses (*53, 91*). While some studies have found that these cross-reactive antibodies can neutralize SARS-CoV-2 *in vitro*, others have not (*61, 91, 92*). In patients, cross-reactive antibodies are also not associated with protection against SARS-CoV-2 infection or severe COVID-19 (*61*). Our findings further suggest that S2-reactive antibodies are not protective, as Tfh-dependent antibodies enriched for S2 epitope reactivity displayed a similar or even lower neutralization potency index against SARS-CoV-2 than more diverse Tfh-independent antibodies.

Our study defines the role of Tfh cells versus LN-Th1 cells in promoting complementary antibody responses during respiratory viral infections. While Tfh cells amplify antibody levels, diversify Fc effector function, and focus the repertoire against immunodominant epitopes, LN-Th1 cells ensure the production of antibodies with activating Fc effector function and broader reactivity. LN-Th1-driven responses may serve as a parallel mechanism for producing protective antibodies in settings of Tfh/GC impairment, such as COVID-19-induced inflammation and old age (*14, 93*). Understanding this additional axis of antiviral antibody production may therefore inform more effective vaccine design and help establish a new paradigm for how T cell-dependent humoral immunity is generated.

## Acknowledgements

We thank all members of the Eisenbarth and Wilen labs for helpful discussions. We would like to acknowledge Benhur Lee, BEI Resources, Joerg Nikolaus, and Yale West Campus Imaging Core for providing critical reagents, resources, and expertise. We thank Yale Environmental Health and Safety for providing necessary training and support for SARS-CoV-2 research. Illustrations were created with BioRender.com.

## Author contributions

Conceptualization: JSC, CBW, SCE

Methodology: ES, TM, BI, KK, JB, WAH, JW, MMA, WS, JSW

Investigation: JSC, ES, TM, KK, JB, RBF, BLM

Validation: JSC, RDC, ES, TM, KK, JB, WAH

Formal Analysis: JSC, RDC, WAH

Visualization: JSC, RDC

Resources: LP, LC, SC, JC, JCS, AI, CBW, SCE

Writing: JSC, RDC, CBW, SCE

Supervision: JSW, UG, CBW, SCE

Funding acquisition: CBW, SCE

## Data and materials availability

All data are available in the main text or the supplementary materials.

## Materials and Methods

### Mice

*Bcl6*^fl/fl^ [B6.129S(FVB)-*Bcl6^tm1.1Dent^*/J (*30*)], *Cd4*^Cre^ [B6.Cg-Tg(Cd4-cre)1Cwi/BfluJ (*94*)], *Ciita*^−/−^ [B6.129S2-*Ciita^tm1Ccum^*/J (*35*)], K18-hACE2 [B6.Cg-Tg(K18-ACE2)2Prlmn/J (*48*)] were purchased from Jackson Laboratory. *Bcl6*^fl/fl^ mice were crossed with *Cd4*^Cre^ mice to generate *Bcl6*^fl/fl^*Cd4*^Cre^ mice. K18-hACE2 mice were crossed with *Bcl6*^fl/fl^*Cd4*^Cre^ mice to generate K18-hACE2 *Bcl6*^fl/fl^ and *Bcl6*^fl/fl^*Cd4*^Cre^ mice. Mice were bred in-house using mating trios to enable utilization of littermates for experiments. Mice of both sexes between 6 and 10 weeks old were used for this study. Animal use and care was approved in agreement with the Yale Animal Resource Center and Institutional Animal Care and Use Committee according to the standards set by the Animal Welfare Act.

### Cell lines

Huh7.5, Vero-E6, and 293T cells were from ATCC. Cell lines were cultured in Dulbecco’s Modified Eagle Medium (DMEM) with 10% heat-inactivated fetal bovine serum (FBS) and 1% Penicillin/Streptomycin. All cell lines tested negative for *Mycoplasma* spp.

### AAV-hACE2 transduction

Adeno-associated virus 9 expressing hACE2 from a CMV promoter (AAV-hACE2) was purchased from Vector Biolabs (SKU AAV-200183). Mice were anesthetized by intraperitoneal injection of ketamine (50 mg/kg) and xylazine (5 mg/kg). The rostral neck was shaved and disinfected with povidone-iodine. After a 5-mm incision was made, the salivary glands were retracted and the trachea visualized. Using a 31G insulin syringe, 10^11^ genomic copies of AAV-hACE2 in 50 μl PBS were injected into the trachea. The incision was closed with 3M Vetbond tissue adhesive, and mice were monitored until full recovery.

### Viruses

SARS-CoV-2 P1 stock was generated by inoculating Huh7.5 cells with SARS-CoV-2 isolate USA-WA1/2020 (BEI Resources, NR-52281) at a MOI of 0.01 for three days. The P1 stock was then used to inoculate Vero-E6 cells, and the supernatant was harvested after three days at 50% cytopathic effect. The supernatant was clarified by centrifugation (450 × *g* for 5 min), filtered through a 0.45-micron filter, and stored in aliquots at -80°C. For infection of AAV-hACE2 mice, virus was concentrated by mixing one volume of cold 4X PEG-it Virus Precipitation Solution (40% wt/vol PEG-8000 and 1.2 M NaCl) with three volumes of viral supernatant. The mixture was incubated overnight at 4°C and then centrifuged at 1500 × *g* for 60 min at 4°C. The pelleted virus was resuspended in PBS and stored in aliquots at -80°C. Virus titer was determined by plaque assay using Vero-E6 cells (*95*).

Influenza virus A/PR/8/34 H1N1 (PR8) expressing the ovalbumin OT-II peptide was grown for 2.5 days at 37°C in the allantoic cavities of 10-day-old specific-pathogen-free fertilized chicken eggs. Harvested virus was centrifuged at 3000 × *g* for 20 min at 4°C to remove debris and stored in aliquots at -80°C. Virus titer was determined by plaque assay on Madin-Darby canine kidney cells (*96*).

For all infections, mice were anesthetized using 30% vol/vol isoflurane diluted in propylene glycol (30% isoflurane) and administered SARS-CoV-2 or PR8 intranasally in 50 μl PBS. AAV-hACE2 *Bcl6*^fl/fl^, *Bcl6*^fl/fl^*Cd4*^Cre^, and *Ciita*^−/−^ mice were infected with 1.2×10^6^ PFU of SARS-CoV-2. K18-hACE2 *Bcl6*^fl/fl^ and *Bcl6*^fl/fl^*Cd4*^Cre^ mice were infected with 20 PFU of SARS-CoV-2. K18-hACE2 mice in serum transfer experiments were infected with 100 PFU of SARS-CoV-2. *Bcl6*^fl/fl^, *Bcl6*^fl/fl^*Cd4*^Cre^, and *Ciita*^−/−^ mice were infected with 70 PFU of PR8. All work with SARS-CoV-2 was performed in a biosafety level 3 (BSL3) facility with approval from the office of Environmental Health and Safety and the Institutional Animal Care and Use Committee at Yale University.

### Enzyme-linked immunosorbent assay (ELISA)

Sera were incubated with a final concentration of 0.5% Triton X-100 and 0.5 mg/ml RNase A for 30 min at room temperature to inactivate potential SARS-CoV-2. SARS-CoV-2 stabilized spike glycoprotein (BEI Resources, NR-53257) (*97*), SARS-CoV-2 spike glycoprotein receptor-binding domain (RBD) (BEI Resources, NR-52946), and influenza virus A/PR/8/34 H1N1 hemagglutinin (HA) protein (Sino Biological, 11684-V08H) were coated at a concentration of 2 μg/ml in carbonate buffer on 96-well MaxiSorp plates (Thermo Fisher) overnight at 4°C. PR8 was inactivated with 1% Triton X-100 for 1 hr at 37°C and coated at a concentration of 20 μg/ml in carbonate buffer. Plates were blocked with 1% BSA in PBS for 1 hour at room temperature. Serum samples were serially diluted in 1% BSA in PBS and incubated in plates for 2 hours at room temperature. Antibody isotypes were detected with anti-mouse IgG-HRP (1013-05), anti-mouse IgG1-HRP (1073-05), anti-mouse IgG2b-HRP (1093-05), anti-mouse IgG2c-HRP (1077-05), or anti-mouse IgG3-HRP (1103-05) from Southern Biotech or anti-mouse IgM-HRP (550588) from BD Biosciences by incubating for 1 hr at room temperature. Plates were developed with TMB Stabilized Chromogen (Thermo Fisher), stopped with 3 N hydrochloric acid, and read at 450 nm on a microplate reader. Background was determined as twice the average OD value of blank wells. Pooled sera from mice infected with SARS-CoV-2 or PR8 were used as reference standards to calculate arbitrary units. To measure antibody affinity, serial dilutions of serum samples were plated in duplicate. After incubation of serum samples, a 10-min wash with 5.3 M urea was performed on one set of the samples. Percentage of IgG binding after urea wash was calculated by dividing the area under the curve for each sample with urea wash by that without urea wash.

### Flow cytometry

Mediastinal lymph nodes (medLN) were homogenized with a syringe plunger and filtered through a 70-μm cell strainer. Red blood cells were lysed with RBC Lysis Buffer (BioLegend) for 1 min. Single-cell preparations were resuspended in PBS with 2% FBS and 1 mM EDTA and pre-incubated with Fc block (clone 2.4G2) for 5 min at room temperature before staining. Cells were stained with the following antibodies or viability dye for 30 min at 4°C: anti-CD4 (RM4-5), TCRβ (H57-597), PD-1 (RMP1-30), CD44 (IM7), PSGL-1 (2PH1), Ly6C (HK1.4), B220 (RA3-6B2), Fas (Jo2), GL7 (GL7), CD138 (281-2), and LIVE/DEAD™ Fixable Aqua (Thermo Fisher). CXCR5 (L138D7) was stained for 25 min at room temperature.

CD40L staining was performed by surface mobilization assay (*98*). Cells were pre-incubated with Fc block (10 μg/ml) for 10 min at room temperature. Cells were then stimulated with 50 ng/ml PMA and 1 μg/ml ionomycin in complete RPMI (10% heat-inactivated FBS, 1% Penicillin/Streptomycin, 2 mM L-glutamine, 1mM sodium pyruvate, 10 mM HEPES, 55 μM 2-mercaptoethanol) in the presence of PE/Cy7 anti-CD40L (MR1, 4 μg/ml) for 30 min at 37°C. After washing, the remaining markers were stained as described above.

For intracellular cytokine staining, cells were stimulated with 50 ng/ml PMA and 500 ng/ml ionomycin in complete IMDM (10% heat-inactivated FBS, 1% Penicillin/Streptomycin, 2 mM L-glutamine, 1mM sodium pyruvate, 25 mM HEPES, 1X MEM Non-Essential Amino Acids, 20 μM 2-mercaptoethanol) in the presence of BD GolgiPlug™ (1:1000) for 2 hr at 37°C. Cells were then washed, incubated with Fc block, and stained for surface markers PD-1, PSGL-1, and Ly6C as well as LIVE/DEAD™ Fixable Aqua. Cells were next fixed with 2% paraformaldehyde in PBS for 30 min at 4°C, followed by permeabilization with 1X eBioscience™ Permeabilization Buffer for 15 min 4°C. Intracellular staining was performed for CD4, CD44, CXCR5, IFN-γ (XMG1.2), and IL-21 (1 μg of IL-21R-Fc Chimera Protein; R&D Systems, 596-MR-100) in permeabilization buffer for 40 min at 4°C, followed by secondary staining with PE anti-human IgG (Jackson ImmunoResearch, 109-116-098) for 30 min at 4°C.

Samples from SARS-CoV-2-infected mice were fixed with 4% paraformaldehyde for 30 min at room temperature before removal from BSL3 facility. Samples were acquired on a CytoFLEX S (Beckman Coulter) and analyzed using FlowJo software (BD).

### Immunofluorescence

MedLN were fixed with 4% paraformaldehyde in PBS for 4 hr at 4°C, followed by cryopreservation with 20% sucrose in PBS for 2 hr at 4°C. MedLN were snap-frozen in optimal cutting temperature compound and stored at -80°C. Tissues were cut into 7-μm sections and blocked with 10% rat serum in staining solution (PBS with 1% BSA and 0.1% Tween-20) for 1 hr at room temperature. Sections were stained with BV421 anti-PSGL-1 (2PH1) and AF647 anti-CD4 (RM4-5) or BV421 anti-B220 (RA3-6B2) and AF647 anti-GL7 (GL7) along with Fc block, biotinylated anti-mouse IgG2c (5.7), and rabbit anti-T-bet (E4I2K; Cell Signaling Technology, 97135S) overnight at 4°C. Secondary staining was performed with AF594 streptavidin and AF488 goat anti-rabbit IgG for 1 hr at room temperature. Images were acquired on a Nikon TiE Spinning Disk Confocal Microscope with the 30× objective and analyzed with ImageJ.

### Pseudovirus production

Vesicular stomatitis virus (VSV)-based pseudotyped viruses were produced as previously described (*95, 99*). Vector pCAGGS containing the SARS-CoV-2 Wuhan-Hu-1 spike glycoprotein gene was produced under HHSN272201400008C and obtained through BEI Resources (NR-52310). The sequence of the Wuhan-Hu-1 isolate spike glycoprotein is identical to that of the USA-WA1/2020 isolate. The spike sequence of the B.1.351 variant of concern was generated by introducing the following mutations: L18F, D80A, D215G, R246I, K417N, E484K, N501Y, and A701V. 293T cells were transfected with either spike plasmid, followed by inoculation with replication-deficient VSV expressing *Renilla* luciferase for 1 hour at 37°C (*99*). The virus inoculum was then removed, and cells were washed with PBS before adding media with anti-VSV-G (8G5F11) to neutralize residual inoculum. Supernatant containing pseudovirus was collected 24 hours post inoculation, clarified by centrifugation, concentrated with Amicon Ultra Centrifugal Filter Units (100 kDa), and stored in aliquots at -80°C. Pseudoviruses were titrated in Huh7.5 cells to achieve a relative light unit (RLU) signal of 600 times the cell-only control background.

### Pseudovirus neutralization assay

Sera for neutralization assay were heat-inactivated for 30 min at 56°C. Sera were tested at a starting dilution of 1:50 for USA-WA1/2020 pseudovirus and 1:12.5 for B.1.351 pseudovirus, with up to eight two-fold serial dilutions. 2×10^4^ Huh7.5 cells were plated in each well of a 96-well plate the day before. Serial dilutions of sera were incubated with pseudovirus for 1 hour at 37°C. Growth media was then aspirated from the cells and replaced with 50 µl of serum/virus mixture. Luciferase activity was measured at 24 hours post infection using the *Renilla* Luciferase Assay System (Promega). Each well of cells was lysed with 50 µl Lysis Buffer, and 15 µl cell lysate was then mixed with 15 µl Luciferase Assay reagent. Luminescence was measured on a microplate reader (BioTek Synergy). Half maximal inhibitory concentration (IC50) was calculated as previously described (*100*). Neutralizing titer (NT50) was defined as the inverse of IC50.

### Serum transfer

Sera from SARS-CoV-2-infected AAV-hACE2 *Bcl6*^fl/fl^ or *Bcl6*^fl/fl^*Cd4*^Cre^ mice at 14 days post infection (dpi) were pooled, and the resulting levels of spike-specific IgG were measured by ELISA. *Bcl6*^fl/fl^ sera was diluted with naïve sera to match the spike-specific IgG titer of *Bcl6*^fl/fl^*Cd4*^Cre^ sera. Serum samples were then diluted 1:1 with PBS, and 200 µl of diluted serum was transferred intravascularly by retro-orbital injection into K18-hACE2 mice under anesthesia with 30% isoflurane.

### Measurement of lung viral burden

The left lobe of the lung was collected and homogenized in 1 ml DMEM supplemented with 2% heat-inactivated FBS and 1% Antibiotic-Antimycotic. Lung homogenates were clarified by centrifugation at 3200 × *g* for 10 min and stored in aliquots at -80°C. Viral burden was measured in lung homogenates by plaque assay on Vero-E6 cells as previously described (*95*). In addition, 250 µl of lung homogenate was mixed with 750 µl of TRIzol™ LS Reagent (Invitrogen), and RNA was extracted with the RNeasy Mini Kit (Qiagen) following manufacturer’s instructions. cDNA synthesis was performed using random hexamers and ImProm-II™ Reverse Transcriptase (Promega). Quantitative PCR (qPCR) was performed in triplicate for samples and standards using SARS-CoV-2 nucleocapsid (N1)-specific oligonucleotides: Probe: 5’ 6FAM-ACCCCGCATTACGTTTGGTGGACC-BHQ1 3’; Forward primer: 5’GACCCCAAAATCAGCGAAAT-3’; Reverse primer: 5’ TCTGGTTACTGCCAGTTGAATCTG 3’. The limit of detection was 100 SARS-CoV-2 genome copies/μl. Virus copy numbers were quantified using a control plasmid containing the complete nucleocapsid gene from SARS-CoV-2. SARS-CoV-2 genome copies were normalized to *Actb* using *Actb*-specific oligonucleotides: Probe: 5’ 6-JOEN-CACCAGTTC /ZEN/ GCCATGGATGACGA-IABkFQ 3’; Forward primer: 5’ GCTCCTTCGTTGCCGGTCCA 3’; Reverse primer: 5’ TTGCACATGCCGGAGCCGTT 3’. The limit of detection was 100 *Actb* copies/μl. Samples with undetectable SARS-CoV-2 genome copies were set at 0.01 relative to *Actb*.

### Serum epitope repertoire analysis (SERA) with SARS-CoV-2 linear epitope library

Serum samples for epitope profiling were inactivated with UV light (250 mJ). The SERA platform for next-generation sequencing (NGS)-based analysis of antibody epitope repertoires has been previously described (*101*). In brief, *Escherichia coli* were engineered with a surface display vector carrying linear peptides derived from the SARS-CoV-2 proteome (GenBank MN908947.3), designed using oligonucleotides (Twist Bioscience) encoding peptides 12 amino acids in length and tiled with 8 amino acids overlapping. Serum samples (0.5 µl each) were then diluted 1:200 in a suspension of PBS and bacteria carrying the surface display library (10^9^ cells per sample with 3×10^5^ fold library representation), and incubated so that antibodies contained in the serum would bind to the peptides on the surface of the bacteria. After incubating with protein A/G magnetic beads and magnetically isolating bacteria that were bound to antibodies contained in the serum, plasmid DNA was purified and PCR amplified for NGS. Unique molecular identifiers (UMIs) were applied during PCR to minimize amplification bias, designed as an 8 base pair semi-random sequence (NNNNNNHH). After preprocessing and read trimming the raw sequencing data, the resulting reads were filtered by utilizing the UMIs to remove PCR duplicates. The filtered UMI data (hereafter referred to as reads) were then aligned to the original reference of linear epitopes derived from SARS-CoV-2 and quantified.

From the raw mapped read counts for each of the 2410 linear epitopes represented in the library, we first calculated the Shannon entropy and Simpson’s diversity index of each sample using the *diversity* function in the *diverse* R package. To calculate the “repertoire focusing index”, we used the formula: 1-(H’/log_2_(R)), where H’ is Shannon entropy and R is richness (*102*), defined here as the number of unique epitopes recognized by a given sample (read count > 0). Statistical differences in these various metrics were assessed by two-tailed unpaired Welch’s t-test, comparing *Bcl6*^fl/fl^ and *Bcl6*^fl/fl^*Cd4*^Cre^ conditions.

For further analysis, we normalized the raw count data using the median ratio approach implemented in DESeq2 (*103, 104*). Differential enrichment analysis was performed using the Wald test in DESeq2, comparing *Bcl6*^fl/fl^ vs *Bcl6*^fl/fl^*Cd4*^Cre^ samples. Multiple hypothesis correction was performed by the Benjamini-Hochberg method, setting a statistical significance threshold of adjusted *p* < 0.05. After identifying differentially enriched epitopes, the normalized counts were log_2_ transformed (hereafter referred to as log_2_ normalized counts) for downstream visualization and analysis.

For converting the log_2_ normalized counts into relative enrichment scores on a sample-by-sample basis, we scaled the log_2_ normalized counts within each sample to *z-*scores. In this manner, a *z-*score of 0 would correspond to epitopes that exhibited an average level of enrichment in a given sample; a positive *z-*score would indicate that an epitope is relatively enriched in a sample, while a negative *z-*score would denote a low-scoring epitope. Where applicable, statistical significance was assessed on the within-sample *z*-scores by two-tailed unpaired Welch’s t-test. For visualization purposes, we also calculated the average *z*-scores in *Bcl6*^fl/fl^ or *Bcl6*^fl/fl^*Cd4*^Cre^ groups.

Pan-human coronavirus (hCoV) conservation scores were calculated through multiple alignment of several hCoV spike sequences using Clustal Omega (*105*). The following hCoV spike sequences were used: HKU1 (UniProt: Q0ZME7), OC43 (P36334), 229E (P15423), NL63 (Q6Q1S2), SARS-CoV (P59594), SARS-CoV-2 (P0DTC2), and MERS-CoV (K9N5Q8). Amino-acid level conservation scores for SARS-CoV-2 spike were extracted through JalView (*106*).

### Statistical analysis

Data analysis was performed using GraphPad Prism 9 unless otherwise indicated. Data were analyzed using Student’s two-tailed, unpaired t-test; Welch’s two-tailed, unpaired t-test; two-tailed Mann–Whitney test; or two-tailed Wilcoxon signed-rank test, as indicated. P < 0.05 was considered statistically significant.

---

**Figure S1:**
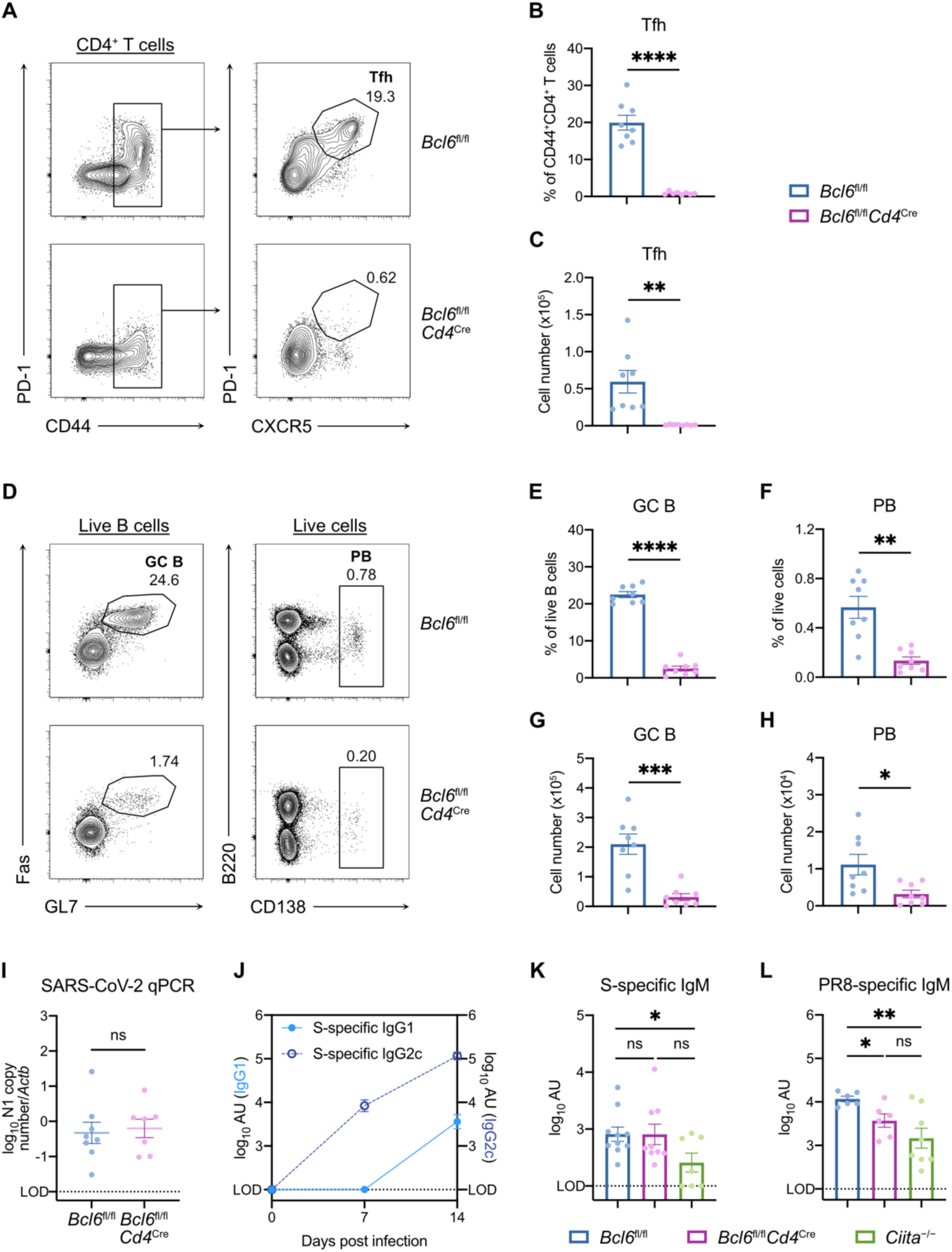
Adaptive immune response to SARS-CoV-2 in Tfh-sufficient and -deficient mice. (**A** to **H**) Flow cytometric analysis of medLN from SARS-CoV-2-infected mice at 14 dpi. (**A**) Representative gating strategy to identify Tfh cells from total CD4^+^ T cells. (**B** and **C**) Frequency among CD44^+^CD4^+^ T cells (B) and total number (C) of Tfh cells in *Bcl6*^fl/fl^ (blue) or *Bcl6*^fl/fl^*Cd4*^Cre^ (magenta) mice. (**D**) Representative gating strategy to identify germinal center (GC) B cells and plasmablasts (PB). (E and F) Frequencies of GC B cells (E) and PB (F) in *Bcl6*^fl/fl^ (blue) or *Bcl6*^fl/fl^*Cd4*^Cre^ (magenta) mice. (**G** and **H**) Total number of GC B cells (G) and PB (H) in *Bcl6*^fl/fl^ (blue) or *Bcl6*^fl/fl^*Cd4*^Cre^ (magenta) mice. (**I**) Viral burden in lungs from *Bcl6*^fl/fl^ mice or *Bcl6*^fl/fl^*Cd4*^Cre^ mice infected with SARS-CoV-2 at 7 dpi, assessed by qPCR. Data are expressed as log_10_ N1 gene copy number by qPCR, normalized to *Actb*. (**J**) Time course of S-specific IgG1 and IgG2c serum antibody titers from *Bcl6*^fl/fl^ mice infected with SARS-CoV-2. (**K**) S-specific IgM titers in sera from *Bcl6*^fl/fl^ (blue), *Bcl6*^fl/fl^*Cd4*^Cre^ (magenta), or *Ciita*^−/−^ (green) mice at 14 dpi with SARS-CoV-2. (**L**) PR8-specific IgM titers in sera from *Bcl6*^fl/fl^ (blue), *Bcl6*^fl/fl^*Cd4*^Cre^ (magenta), or *Ciita*^−/−^ (green) mice at 14 dpi with PR8. Statistical significance was assessed by either two-tailed unpaired t-test or Welch’s t-test, based on the F test for unequal variance. \**P* < 0.05; \*\**P* < 0.01; \*\*\**P* < 0.001; \*\*\*\**P* < 0.0001. ns, not significant. Data are expressed as mean ± standard error of mean (SEM). Each symbol in (B and C, E to I, K and L) represents an individual mouse. Each symbol in (J) represents the mean of six mice. Data are aggregated from at least two independent experiments.

**Figure S2:**
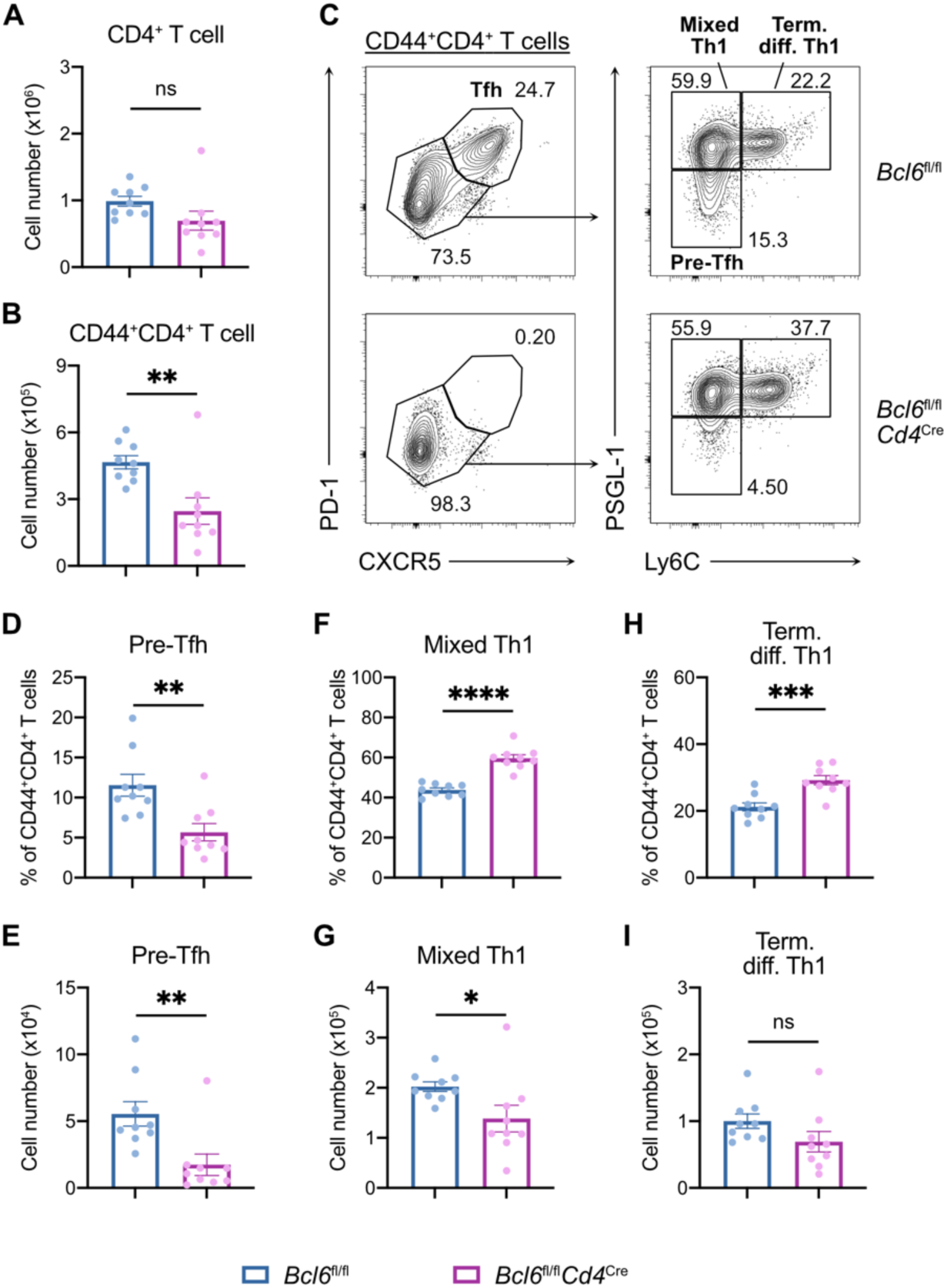
Characterization of CD4^+^ T cell subsets induced by SARS-CoV-2 infection. (**A** to **I**) Flow cytometric analysis of medLN from SARS-CoV-2-infected mice at 7 dpi. (**A** and **B**) Total number of CD4^+^ T cells (**A**) and CD44^+^CD4^+^ T cells (**B**) in *Bcl6*^fl/fl^ (blue) or *Bcl6*^fl/fl^*Cd4*^Cre^ (magenta) mice. (**C**) Representative gating strategy to define Tfh cells, pre-Tfh cells, mixed Th1 cells, and terminally differentiated Th1 cells from CD44^+^CD4^+^ T cells. (**D** to **I**) Frequency among CD44^+^CD4^+^ T cells and total number of pre-Tfh cells (D and E), mixed Th1 cells (F and G), and terminally differentiated Th1 cells (H and I) in *Bcl6*^fl/fl^ (blue) or *Bcl6*^fl/fl^*Cd4*^Cre^ (magenta) mice. Statistical significance was assessed by either two-tailed unpaired t-test or Welch’s t-test, based on the F test for unequal variance. \**P* < 0.05; \*\**P* < 0.01; \*\*\**P* < 0.001; \*\*\*\**P* < 0.0001. ns, not significant. Data are expressed as mean ± SEM. Each symbol represents an individual mouse. Data are aggregated from three independent experiments.

**Figure S3:**
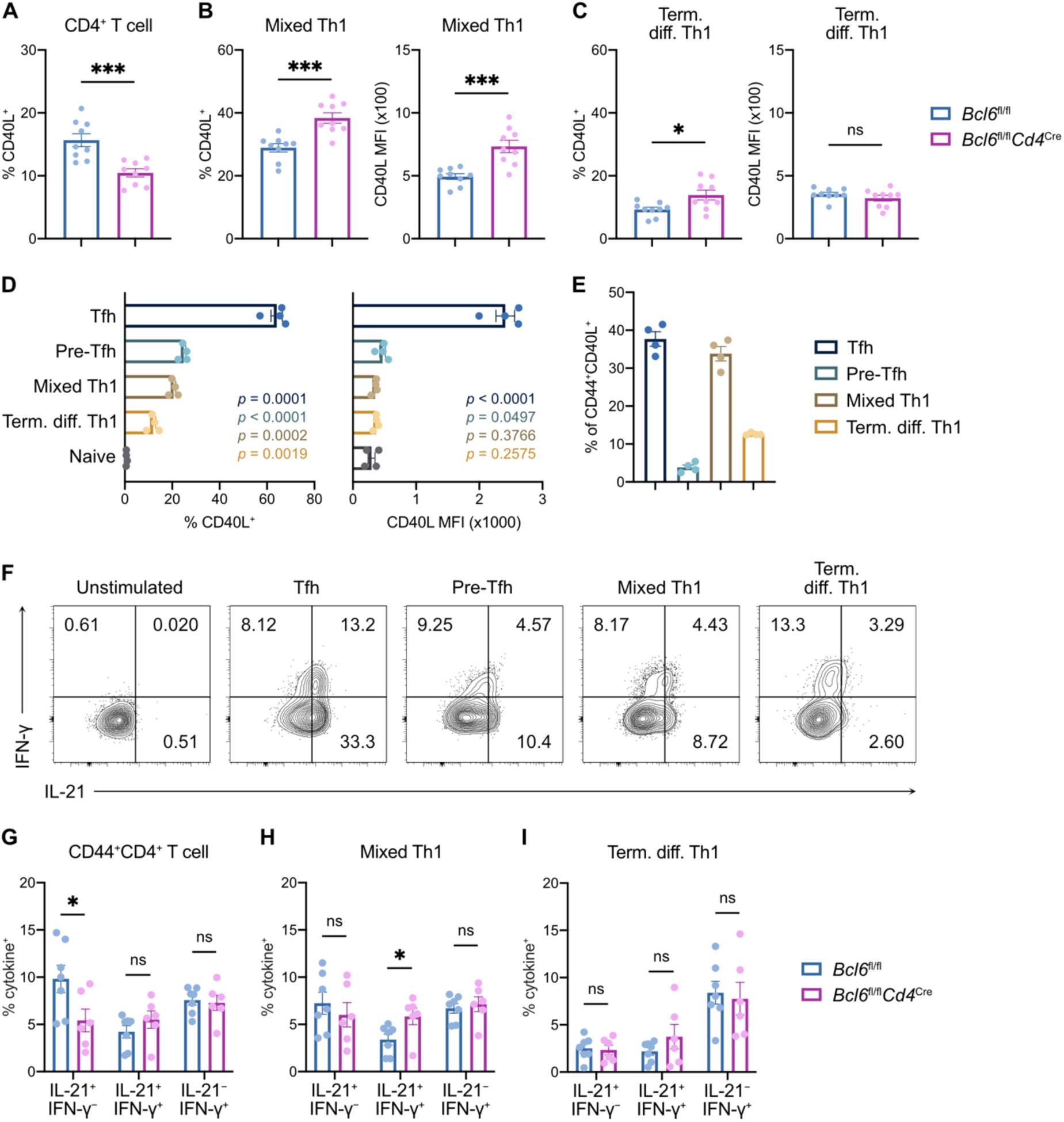
CD40L and cytokine expression in CD4^+^ T cell subsets after viral infection. (**A** to **C**) Flow cytometric analysis of medLN from *Bcl6*^fl/fl^ (blue) or *Bcl6*^fl/fl^*Cd4*^Cre^ (magenta) mice at 7 dpi with SARS-CoV-2. (**A**) Frequency of CD44^+^CD40L^+^ cells among CD4^+^ T cells. (**B** and **C**) Frequency (left) and MFI (right) of CD40L expression among mixed Th1 cells (B) and terminally differentiated Th1 cells (C). (**D** and **E**) Flow cytometric analysis of medLN from *Bcl6*^fl/fl^ mice at 7 dpi with PR8. (**D**) Frequency (left) and MFI (right) of CD40L expression among CD4^+^ T cell subsets. *P* values for each subset relative to naive CD4^+^ T cells are color-coded. (**E**) Relative proportions of CD4^+^ T cell subsets among CD44^+^CD40L^+^ cells. (**F**) Representative flow cytometric analysis of IL-21 and IFN-γ protein expression in distinct CD4^+^ T cell subsets from medLN of SARS-CoV-2-infected mice at 7 dpi. (**G** to **I**) Flow cytometric analysis of medLN from *Bcl6*^fl/fl^ (blue) or *Bcl6*^fl/fl^*Cd4*^Cre^ (magenta) mice at 7 dpi with SARS-CoV-2. Frequencies of IL-21^+^IFN-γ^−^, IL-21^+^IFN-γ^+^, and IL-21^−^IFN-γ^+^ expression among CD44^+^CD4^+^ T cells (G), mixed Th1 cells (H) and terminally differentiated Th1 cells (I). Statistical significance was assessed by either two-tailed unpaired t-test or Welch’s t-test, based on the F test for unequal variance. \**P* < 0.05; \*\*\**P* < 0.001. ns, not significant. Data are expressed as mean ± SEM. Each symbol represents an individual mouse. Data in (A to C and G to I) are aggregated from at least two independent experiments.

**Figure S4:**
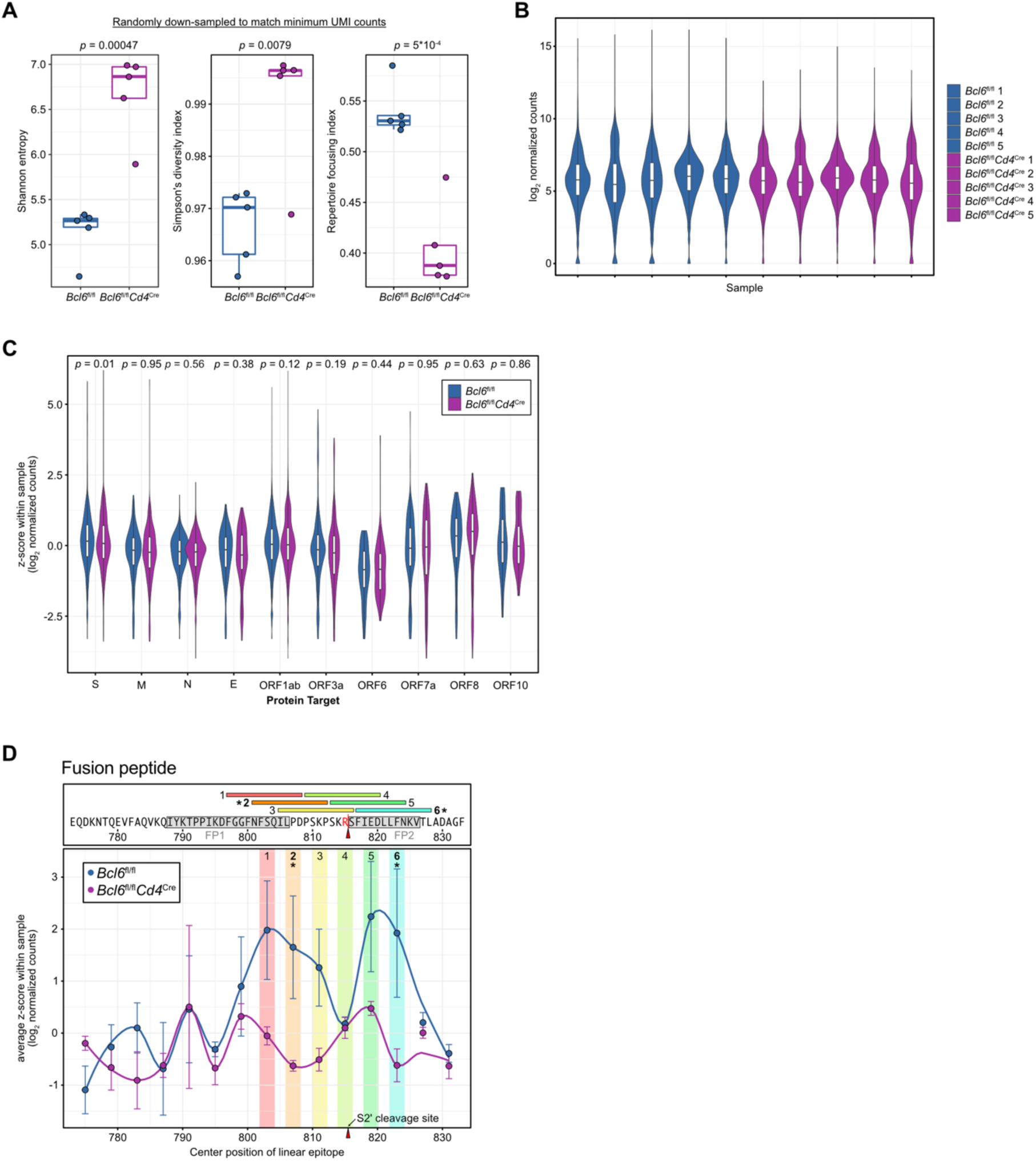
Analysis of SARS-CoV-2 linear epitopes recognized by antibodies from *Bcl6*^fl/fl^ or *Bcl6*^fl/fl^*Cd4*^Cre^ mice. (**A**) Boxplots detailing Shannon entropy (left), Simpson’s diversity index (middle), and repertoire focusing index (right), after random down-sampling to match the sample with the lowest UMI counts. Statistical significance was assessed by two-tailed unpaired Welch’s t-test. Each symbol represents an individual mouse. (**B**) Violin plot of log_2_ normalized counts for all 2410 epitopes in each sample. Higher values indicate relative enrichment of antibodies that are reactive against a particular epitope. (**C**) Violin plot detailing the relative enrichment of antibodies against different SARS-CoV-2 proteins, such that linear epitopes derived from the same protein are grouped together. Data are expressed as *z*-scores, scaled from log_2_ normalized counts within each sample; data are aggregated together by experimental group. Statistical significance was assessed by two-tailed unpaired Welch’s t-test. (**D**) Linear epitopes surrounding the S2’ site (R815, red arrowhead), cleavage of which triggers fusion, are color-coded and numbered with their corresponding amino acid sequences annotated above. Epitopes that are differentially enriched in *Bcl6*^fl/fl^ vs *Bcl6*^fl/fl^*Cd4*^Cre^ at a library-wide scale (from Figure 4C) are bolded and labeled with an asterisk. Putative fusion peptides are labeled in gray boxes above (FP1, 788-806; FP2, 816-826). Data are expressed as average *z-*scores in *Bcl6*^fl/fl^ or *Bcl6*^fl/fl^*Cd4*^Cre^ mice, with SEM error bars and loess regression lines.

